# GSK-3 regulates CD4-CD8 cooperation needed to generate super-armed CD8+ cytolytic T cells against tumors

**DOI:** 10.1101/2025.03.08.642085

**Authors:** Bastien Moës, Janna Krueger, Alexandra Kazanova, Chen Liu, Yunfeng Gao, Nikhil Anto Ponnoor, Linda Castoun-Puckett, Andres Carlos Oroya Lazo, Lan Huong, Auryane Laure Cabald, Thai Hien Tu, Christopher E. Rudd

**Affiliations:** Department of Medicine and Sciences biomédicales, Université de Montréal, Montréal, QC H3C 3J7, Canada; Dept de microbiologie, infectiologie et immunologie, Université de Montréal; Cell Signaling and Immunotherapy Section, Research Center Hopital Maisonneuve-Rosemont (CR-HMR) Montreal, Quebec H1T 2M4; Department of Medicine, McGill University; University Institute for Hematology-Oncology and Cell Therapy, Hopital Maisonneuve-Rosemont, Montreal, QC H1T 2M4, Canada

## Abstract

While immune checkpoint blockade (ICB) has revolutionized cancer treatment, the key T-cell signaling pathways responsible for its potency remain unclear. GSK-3 is an inhibitory kinase that is most active in resting T-cells. In this study, we demonstrate that GSK-3 facilitates PD-1 blockade, an effect seen by modulating CD4 T-cell help for CD8+ CTL responses against ICB resistant tumors. We show that GSK-3 controls metabolic reprogramming towards glycolysis and synergizes with PD-1 to induce a transcriptional program that reduces suppressive CD4+ Treg numbers while generating super-armed effector-memory CD8+ CTLs that express an unprecedented 7/9 granzymes from the genome. Crucially, we found that GSK-3 cooperates with PD-1 blockade to determine the dependency of CD8+ CTLs on help from CD4+ T-cells. Our study unravels a novel cooperative PD-1 blockade-dependent signaling pathway that potentiates CTL responses against tumors, offering a new strategy to overcome immunotherapy resistance by modulating CD4+ helper and CD8+ cytotoxic functions.

**Significance:** This study demonstrates for the first time that GSK-3 controls the crosstalk between CD4+ and CD8+ T cells, synergizing with anti-PD-1 therapy to overcome resistance to checkpoint blockade and to generate super-armed CD8+ effector cells in cancer immunotherapy. This newly uncovered GSK-3-dependent CD4-CD8 T-cell crosstalk mechanism presents a new approach to enhance anti-PD-1 immunotherapy.

## Introduction

Antigen presenting cells, helper, suppressor T-cells as well as memory and effector subsets orchestrate the nature and amplitude of the adaptive immune response(1). CD4+ and CD8+ T-cells work together to eliminate virally infected cells as well as cancer (2, 3). CD4+ T-cells provide essential help for the development of long-lasting memory and cytotoxic effector CD8 cells (4-6). Without this support, CD8+ T-cell function is suboptimal, impairing their ability to control infections and tumor growth (4, 5, 7, 8). CD4 help can be through various mechanisms that include the production of cytokines such as IL-2, IFN-γ, and TNF-α, and by enhancing dendritic cell function (6). By contrast, CD4+FoxP3+ regulatory T-cells (Tregs) play a crucial role in maintaining immune tolerance while suppressing anti-tumor immune responses (9). The manipulation of CD4+ T-cell help versus Treg activity is a promising strategy for cancer immunotherapy (10, 11).

The continuous long-term exposure to antigens can also cause exhaustion which impairs the clearance of pathogens and tumors (12-14). While the transcription factor, T cell factor 1 (TCF-1) defines early progenitor and memory T cells (15-18), the expression of inhibitory co-receptors such as cytotoxic T-lymphocyte antigen 4 (CTLA-4), programmed death-1 (PD-1), and lymphocyte-activation gene 3 (LAG-3) limit T-cell responses(19). In this context, immune checkpoint blockade (ICB) has revolutionized cancer treatment (20-25). This is mediated by antibodies that block the signaling of inhibitory receptors, while also inducing internalisation (20-25). TCF-1+ T-cells are also essential for successful ICB (26-28), as is the development of effector cytolytic T-cells (CTLs) which release cytotoxic granules that contain perforin and granzymes (GZMs) needed to induce apoptosis in target cells (29, 30). Although granzyme B (GZMB) is the best studied GZM, 9 other GZMs are expressed in mice that act on overlapping and unique protein substrates (31, 32).

In addition, activation signals are needed to elicit the development of effective anti-tumor responses (33, 34). These signals are mediated by protein-tyrosine, serine/threonine and lipid kinases as well as GTP binding proteins and downstream substrates (34-37). In this context, the protein-tyrosine kinase p56^lck^, either free or associated with the CD4 or CD8 co-receptors initiates the activation cascade (33, 38, 39). The kinase phosphorylates T-cell receptor (TCR) associated CD3 and ζ chain immunoreceptor tyrosine-based activation motifs (ITAMs) (33, 38). This then leads to the recruitment of a second zeta-chain-associated protein kinase 70 (ZAP-70) (40). Together, these kinases phosphorylate and activate the function of downstream enzymes and adaptor proteins that are needed to integrate signals for transcription factor entry into the nucleus and gene transcription (41).

Unlike p56^lck^ and ZAP-70 which are activated by receptors on the surface of T-cells, glycogen synthase kinase-3 (GSK-3) is a serine/threonine kinase which is constitutively active in quiescent T-cells and becomes inhibited in response to receptor ligation (42, 43). In mammals, two isoforms of GSK-3 have been identified the 51 kDa GSK-3α isoform and a 47 kDa GSK-3β isoform which are encoded by separate genes (43). The phosphorylation of specific serine residues (Ser9: GSK-3β, Ser21: GSK-3α) inactivate the kinase (44, 45). By contrast, phosphorylation at Tyr216: GSK-3β, Tyr279: GSK-3α enhances activity (46, 47). Both isoforms are expressed in numerous tissues (48) and regulate diverse processes (46, 49). This includes the inhibition of glycogen synthase (50) regulation of β-catenin expression (51-53) as well as T-cell motility (54). In the context of tumor immunity, GSK-3 inactivation can downregulate PD-1 and LAG-3 transcription and expression (42, 55) and in the process, limit tumor growth (42, 54-59). In this regard, the FDA has approved studies using GSK-3 inhibitors to treat various cancers (60, 61).

While immune checkpoint blockade has revolutionized cancer treatment, the key T-cell signaling pathways responsible for its potency remain unclear. Further, the signaling mechanisms that are responsible for CD4-CD8 cooperation in checkpoint blockade therapy have remained elusive. Here, we show for the first time that GSK-3 controls CD4-CD8 T-cell cooperation which operates in synergy with anti-PD-1 to generate super-armed CD8+ effectors in immunotherapy. The genetic downregulation of GSK-3 promoted the presence of progenitor and memory CD8+ TILs while decreasing numbers of suppressor Treg TILs. Further, GSK-3 downregulation, combined with anti-PD-1, unlocked an unprecedented cytotoxic potential in CD8+ T-cells, enabling the expression of a 7 out of 9 granzymes from the mouse genome, a phenomenon not before observed. This uncovered GSK-3-regulated CD4-CD8 T-cell crosstalk mechanism could be exploited to enhance anti-PD-1 immunotherapy.

## Results

### GSK-3 KD mice show increased T-cell memory and less exhaustion in response to Cl 13 LCMV

To define further the role of GSK-3 in T-cell immunity, we first generated mice lacking both GSK-3α and GSK-3β specifically in mature T cells. This was achieved by crossing distal promoter lck-Cre mice (dlck-Cre) with mice carrying floxed alleles for gsk-3α (gsk-3αlox/lox) and gsk-3β (gsk-3βlox/lox). The distal lck promoter drives Cre recombinase expression in peripheral T cells, allowing for gene deletion specifically in mature T cells after their development in the thymus (62). Immunoblotting with an anti-GSK-3α/β antibody showed a >75% reduction in the expression of GSK-3α and GSK-3β in splenic T cells (**Fig. 1A**) (referred to as GSK-3 knock-down (GSK-3KD) or dLCK-CRE).

**Fig. 1:**
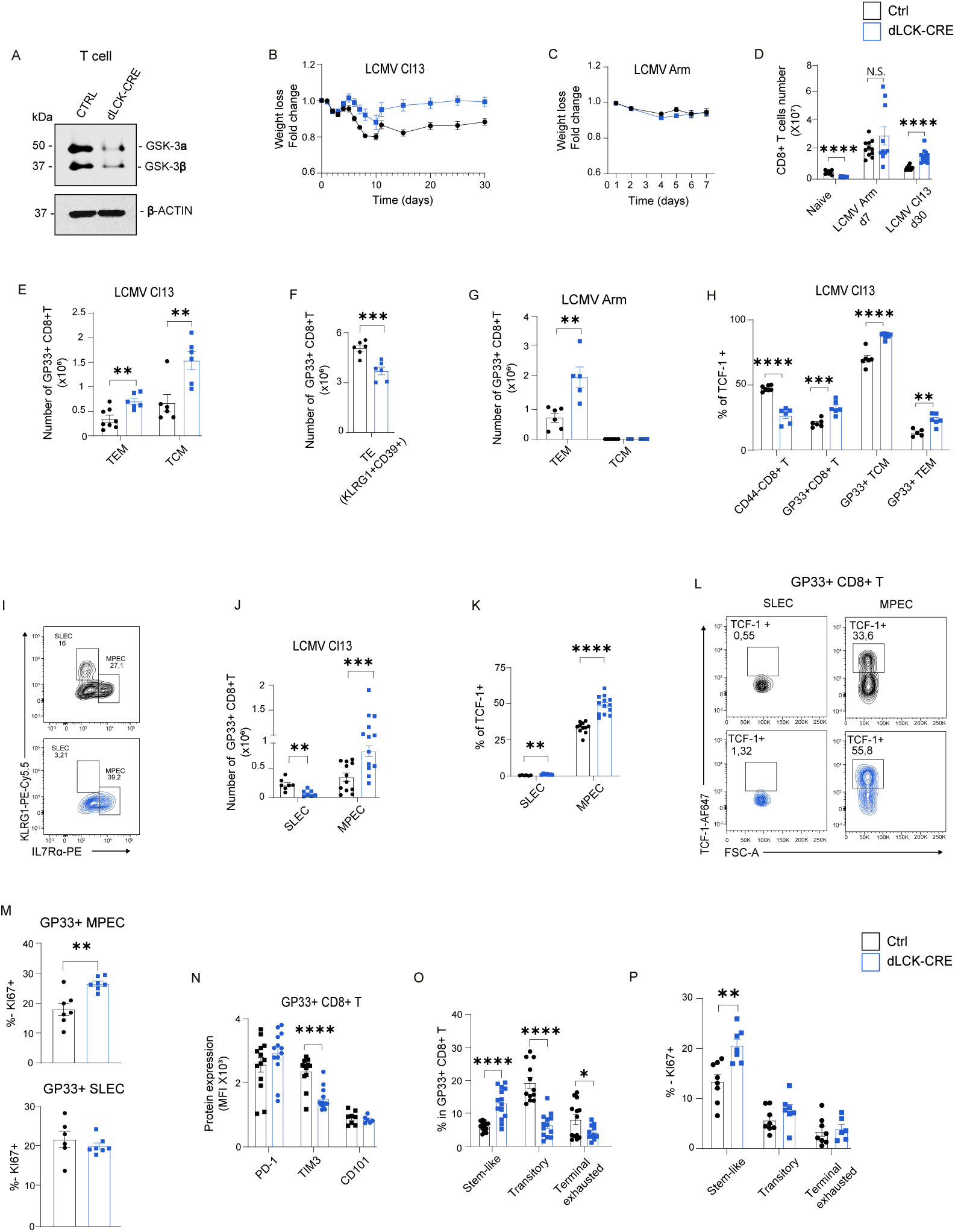
GSK-3 KD regulates T-cell progenitors and exhaustion. (**A**) Western blot analysis of GSK-3 expression in splenic T cells (n=3) Characterization of CD8+ T cell responses following LCMV infection. Mice were infected with LCMV clone 13 (cl13) or Armstrong strain, and spleens were harvested at day 30 or day 7 post-infection (p.i.), respectively. (**B-C**) Weight loss following infection with (**B**) LCMV-cl13 (n=14, pooled from 3 independent experiments) or (**C**) LCMV Armstrong (n=5, representative of 2 independent experiments). (**D**) Splenic CD8+ T cell counts in naive and LCMV-infected mice (Armstrong d7 p.i. and CL-13 d30 p.i.; n=6-14, pooled from ≥2 independent experiments). (**E**) Flow cytometry analysis and quantification of GP33+ CD8+ effector memory (TEM: CD62L-CD44+) and central memory (TCM: CD44+CD62L+) T cells (n=6-8, pooled from 2 independent experiments). (**F**) Quantification of GP33+ CD8+ terminal effector (KLRG1+CD39+) cells (n=6, pooled from 2 independent experiments). (**G**) Proportion of TEM and TCM among GP33+ CD8+ T cells after LCMV Armstrong infection (n=5-6, pooled from 2 independent experiments). (**H**) The precent TCF-1 expression in naive, activated GP33+, TCM, and TEM CD8+ T cells (n=5-7, representative of 2 independent experiments). (**I-J**) Flow cytometry profile (**I**) and histogram quantification (**J**) of short-lived effector cells (SLEC: KLRG1+CD127-) and memory precursor effector cells (MPEC: KLRG1-CD127+) among GP33+ CD8+ T cells (n=7-12, pooled from 2 independent experiments). (**K-L**) Percent TCF-1 expression (**K**) and representative plots (**L**) in SLEC and MPEC populations (n=7-12, pooled from 2 independent experiments). (**M**) Histogram showing Ki67 expression in SLEC and MPEC populations (n=6-7). Upper panel: GP33+MPEC+; lower panel: GP33+SLEC+ (**N**) Expression of exhaustion markers PD1, TIM3, and CD101 on GP33+ CD8+ T cells (n=12-14, pooled from 2 independent experiments). (**O**) Distribution of stem-like (TIM3-TCF1+), transitory (CX3CR1+TIM3+CD101- ), and terminal exhausted (CX3CR1-TIM3+CD101+) cells among PD1+GP33+ CD8+ T cells (n=12-14, pooled from ≥2 independent experiments). (**P**) Ki67 expression in stem-like, transitory, and terminal exhausted populations (n=7-8). Data presented as mean ± SEM. Statistical significance determined by unpaired t-test: *p<0.05, **p<0.01, ***p<0.001, ****p<0.0001; NS, not significant.

We next assessed whether GSK-3 KD would affect responses to infection by chronic Cl 13 and acute Armstrong stains of LCMV, as described (42, 63). In the case of Cl13, the dLck-CRE GSK-3 KD mice showed less weight loss from days 0-8 followed by a more rapid and complete recovery from day 10-30, By contrast, wild type mice showed a more severe loss of weight and a delayed recovery (**Fig. 1B**). Even by 30 days the weight of wild type mice had not fully recovered to the level seen in dLck-CRE mice. By contrast, as a control, no difference in the weight was observed following acute infection with LCMV Armstrong (**Fig. 1C**).

In terms of virally reactive cells, the spleens of GSK-3KD mice had higher levels of CD8+ T-cells than wild-type mice at day 30 in response to Cl13 and at day 7 in response to the Armstrong strain (**Fig. 1D**). Among these cells, GSK-3KD mice showed greater numbers of gp33 reactive effector memory (GP33+CD44+CD62L-) and central memory (GP33+CD44+CD62L+) CD8+ T-cells in response to Cl13 as detected with tetramers containing the GP33 peptide (**Fig. 1E**). Conversely, the numbers of GP33+ KLRG1+CD39+ terminal effector cells (TE) were reduced in GSK-3KD mice (**Fig. 1F**). This increased generation of dLCK-CRE CD8 effector memory was also seen in response to the Armstrong strain. Central memory cells were not detectable at this earlier 7-day time point (**Fig. 1G**). These data demonstrate that reduced GSK-3 promotes the differentiation of CD8+ T-cells into effector memory and central memory T-cells.

In terms of other subsets, we also observed increased numbers of CD4+ T-cells, CD44+CD4+ T-cells, and CD44+Gp33+CD4+ T-cells in GSK-3KD mice relative to controls as well as Th1, Th2 and FoxP3 Tregs (**Supplementary Figure S1**). In addition, we observed an increase in total B cells, CD44+ B cells, and CD44+GP33+ B cells in GSK-3KD mice compared to controls as well as germinal center CD44+ B-cell numbers in GSK-3KD mice (**Supplementary Figure S2**).

Within these cells, GSK-3KD mice also showed an increase in the presence of the progenitor CD8+ T-cell marker TCF1 (**Fig. 1H**). While the expression of TCF1 in the CD8+CD44-population was lower in the GSK-3KD mice, the percent representation of GP33+TCF1+ CD8+ T-cells within the CD44+ pool of experienced T-cells was routinely higher in the GSK-3KD mice (**Fig. 1H**). This was observed in the overall CD8+GP33+ population as well as in the central and effector memory CD8+ compartments.

Consistent with these findings, GSK-3 KD mice gave rise to greater numbers of memory precursor effector cells (MPECs) relative to short-lived effector cells (SLECs) when compared to WT mice (**Fig. 1I**). MPECs were defined as KLRG1-IL7Rα+ while SLECs were defined as KLRG1+IL7Rα-. A FACs plot of a representative experiment showed an increase in MPECs (i.e., 27.1 to 39.2) in GSK-3 KD compared to WT mice. Conversely, there was a corresponding decrease in SLECs. In a comparison of multiple mice, a similar effect was observed where the numbers of GP33+CD8+ MPEC was higher in the GSK-3KD mice than in the WT mice (**Fig. 1J**).

Interestingly, TCF1 was selectively expressed in the MPEC population, and this expression was higher in the GSK-3KD CD8+ cells (**Fig. 1K**) and in a presentative FACs plot of GP33+CD8+ T-cells from 33.6% in WT to 55.8% in GSK-3KD MPECs (**Fig. 1L**). Further, importantly, cell division was only seen in the GP33+ MPEC population as shown by the increase in the cell cycle marker Ki67 (**Fig. 1M**, upper panel). Ki67 is marker for cell division(64). The shorter lived SLECs also showed the presence of dividing cells but they were similar in WT and KD mice (**Fig. 1M**, lower panel).

The reduction in GSK-3 also affected stages of CD8+ T-cell exhaustion. The stages include progenitor PD-1+TCF-1+TIM3-cells which develop into transitionary cells (PD-1+TIM-3+CX3CR1+) and finally into terminally exhausted (PD-1+TIM-3+CX3CR1-CD101+) cells (65, 66). In this context, the expression of TIM-3 was significantly reduced in GSK-3 KD GP33+ CD8+ T-cells (**Fig. 1N**). Exhausted T-cells are characterized by high levels of TIM-3 with reduced effector functions such as decreased cytokine production and cytotoxicity (65, 66). Overall, in terms of GP33+ PD-1+ T-cells, we observed an increase in stem-like TCF-1+ progenitors concurrent with a reduction in both transitory and terminal exhausted CD8+ T-cells (**Fig. 1O**). The same TCF-1+ progenitors also showed an increase in cell division as shown by Ki67 expression (**Fig. 1P**). Similar results were seen in an *in vitro* model of exhaustion where splenic T cells were restimulated over 13 days (**Supplementary Figure S3**). The GSK-3-/- Lck Cre T cells showed reduced levels of PD-1 and TIM-3 expression (**Fig S3B**) and concurrently, a frequency of IFN-γ and TNF-α co-producing cells (**Fig S3C**). Overall, these data show for the first time using a genetic knock-down model that GSK-3 plays a central role in the generation of stem-like progenitors in the process that limits T-cell exhaustion.

### GSK-3KD induces metabolic reprogramming toward glycolysis in T-cells

The increased presence of CD8+ progenitor and memory cells in dLCK-CRE infected mice suggested that the mediator might control the sensitivity of T-cells to TCR ligation. To assess this, isolated CD8+ T-cells were labelled with Tag-it and incubated in vitro with a titration of soluble anti-CD3 antibodies from a range of 0.005 to 2ug/ml over 48-62 hours (**Fig. 2A**). Under these conditions, the proliferative response between KD and WT T-cells were similar. The one exception was the response to lowest concentration of 0.005ug/ml anti-CD3 where the KD T-cells exhibited a clearly better response (see arrow). A similar result was obtained when we stained for Ki67, a marker for cell division (**Fig. 2B**). This clearly demonstrated that GSK-3 controls the sensitivity of T-cells to a remarkably low concentration of anti-CD3. These data suggest that GSK-3 might eventually be found to protect against responses to low affinity or abundance self-antigen.

**Fig. 2:**
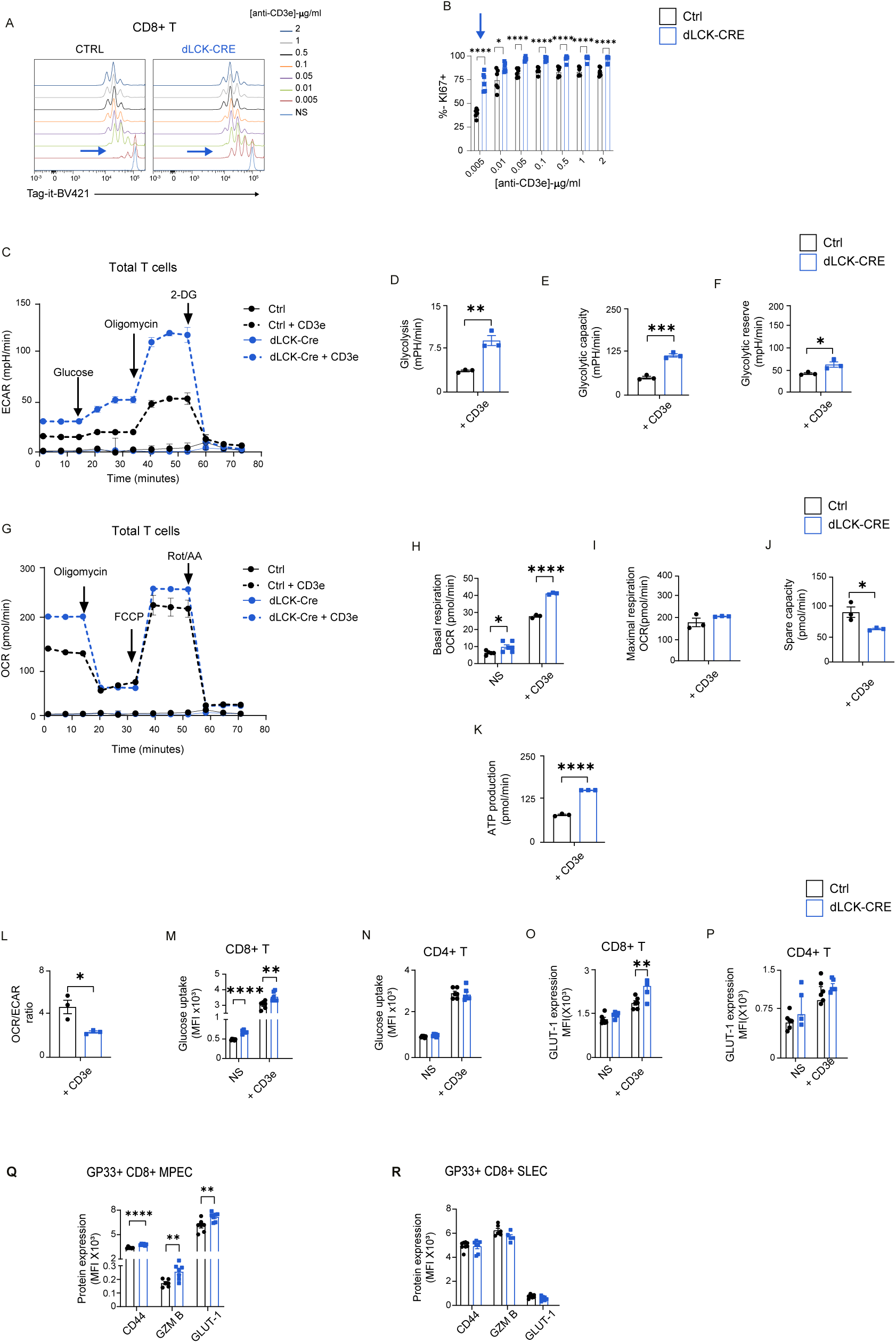
Metabolic profiling of T cells from WT and dLCK-CRE mice. (**A**) Representative flow cytometry plots of Tag-it staining in splenic CD8+ T cells stimulated with varying concentrations of anti-CD3ε monoclonal antibodies (mAbs) for 48 hours. (**B**) Proportion of KI67+ cells among CD8+ T cells, with blue arrow indicating lowest anti-CD3ε mAb concentration tested (n=6, pooled from 2 independent experiments). (**C**) Extracellular acidification rate (ECAR) in splenic T cells following 24h activation with anti-CD3ε (1μg/mL). Sequential injections: glucose, oligomycin, and 2-DG. (**D-F**) Quantification of glycolytic parameters: glycolysis (**D**), glycolytic capacity (**E**), and glycolytic reserve (**F**) (n=3). (**G**) Oxygen consumption rate (OCR) in T cells activated with anti-CD3ε (1μg/mL) for 24h. Sequential injections: oligomycin, FCCP, and rotenone/antimycin A (Rot/AA). (**G-K**) Quantification of respiratory parameters: basal respiration, maximal respiration, spare capacity, and ATP production (n=3). (**L**) Ratio of maximal OCR to maximal ECAR. (**M-N**) Glucose uptake measured by NDBG mean fluorescence intensity (MFI) in CD8+ and CD4+ T-cells, unstimulated (NS) or activated with anti-CD3ε (1μg/mL) for 24h (n=5-6). (**O-P**) GLUT-1 expression analysis by flow cytometry in CD8+ and CD4+ T cells, unstimulated (NS) or activated with anti-CD3ε (1μg/mL) for 24h (n=5-6). Data represent mean ± SEM from 2-3 independent experiments. (**Q-R**) GLUT-1, CD44 and GZMB MFI expression analysis by flow cytometry in GP33+CD8+ MPEC+ (O) and GP33+CD8+ SLEC+ T cells (P) from LCMVCl13 infected mice (see Figure 2). Statistical analysis: unpaired t-test (*p<0.05, **p<0.01, ***p<0.001, ****p<0.0001; NS, not significant).

T-cell activation involves a switch from oxidative phosphorylation to glycolysis (67-70). The role of GSK-3 in T-cells has not been examined. We therefore measured oxygen consumption rate (OCR) and extracellular acidification rate (ECAR) in resting and anti-CD3 stimulated T-cells at 24 hours using Seahorse XF24 Extracellular Flux Analyzer (**Fig. 2C-K**). Cells were subjected to the sequential addition of mitochondria perturbing agents such as oligomycin, to block ATP synthesis, FCCP to uncouple the electron transport chain, and rotenone and antimycin A which block ETC complex I and III, respectively). ECAR as a measure of glycolysis as shown in a representative experiment (**Fig. 2C**). Both basal and anti-CD3 induced glycolysis with the addition of exogenous glucose was clearly higher GSK-3 KD T-cells. Comparative measurements from different mice confirmed a significance in glycolysis in response to anti-CD3 (**Fig. 2D**). In addition, the glycolytic capacity in the GSK-3KD T cells was higher (**Fig. 2E**) and the glycolitic reserve (i.e., as indicated by the difference between the maximal ECAR after glucose and oligomycin addition and maximal ECAR after glucose injection only)(**Fig. 2F**). These data showed that the reduction of GSK-3 expression augmented glycolysis as induced by anti-CD3.

GSK-3 levels also affected mitochondrial respiration (i.e., OXPHOS) levels as assessed by the OCR response (**Fig. 2G-K**). Basal OCR levels was higher in GSK-3 KD T-cells (**Fig. 2G, H**) as was the response to FCCP (**Fig. 2G**). The addition of FCCP determine the maximal uncoupled respiration rate, which is the maximum rate of oxygen consumption that can be achieved when the mitochondrial respiration is completely uncoupled from ATP synthesis. This is an indicator of the cell’s ability to meet increased energy demands under stress condition. Basal and anti-CD3 induced OCR was higher the GSK-3 KD cells (**Fig. 2G**), while despite a trend, no statiscal difference in the maximal respiration OCR between GSK-3 KD and WT cells was seen (**Fig. 2H**). Lastly, the spare respiration capacity,as indicated by the difference between the maximal OCR (after FCCP injection) and basal OCR was actually decreased in anti-CD3e cells from GSK-3 KD mice (**Fig. 2J**). In keeping with this, an increase of ATP production but a reduction in the OCR/ECAR ratio was observed in T-cells from GSK-3 KD mice (**Fig. 2K)**.

In keeping with these findings, the OCR to ECAR ratio was lower in KD cells (**Fig. 2L)**. Further, the glucose uptake in CD8, but not CD4 cells, was higher in KD T-cells (**Fig 2M**) as was the expression of the glucose transporter-1 receptor (GLUT-1) (**Fig 2O).** GLUT-1 plays a crucial role in facilitating glucose uptake in activated T-cells (71-73). Notably, this increased expression was more pronounced in CD8+ T-cells (**Figs 2O and P**). Further, the elevated GLUT-1 was observed in GP33+ CD8+ MPECs (**Fig 2Q**), but not in SLECs (**Fig. 2R**). Overall, our findings indicated that GSK-3 affected both glycolysis and OXPHOS pathways in cellular metabolism with a particularly strong impace on anti-CD3 glycolysis associated with the MPEC subset of effector memory T-cells.

### GSK-3 empowers anti-PD1 to overcome ICB tumor resistance

Given these combined effects, we next assessed the role of GSK-3 in regulating CD4+CD8+ T-cell cooperation, where CD4+ T cells provide help needed for CD8+ T cells^57^. For this, we derived and used a B16-F10 melanoma subclone (B16-F10 R1) which is resistance to anti-PD-1 treatment. B16-F10 R1 cells were injected intradermally followed by intra-peritoneal injections of SB415286 (GSK-3 SMI), anti-PD1 or SB415286/anti-PD1 every 4 days (**Fig. 3A**). Neither anti-PD-1 nor SB415286 monotherapy affected tumor growth (**Fig. 3B**). However, the combination of anti-PD-1 and SB415286 synergized to reduce tumor growth from days 8-17 based on tumor volume (**Fig. 3B**) and tumor weight when measured at day 15 (**Fig 3C**). Spider-graphs show reduced volume growth of individual tumors in mice (**Fig. 3D**, left panels). In addition, 65% of mice bearing B16-F10 R1 tumors became responsive to combination therapy as opposed to non with monotherapy (**Fig. 3D**, right panel).

**Fig. 3:**
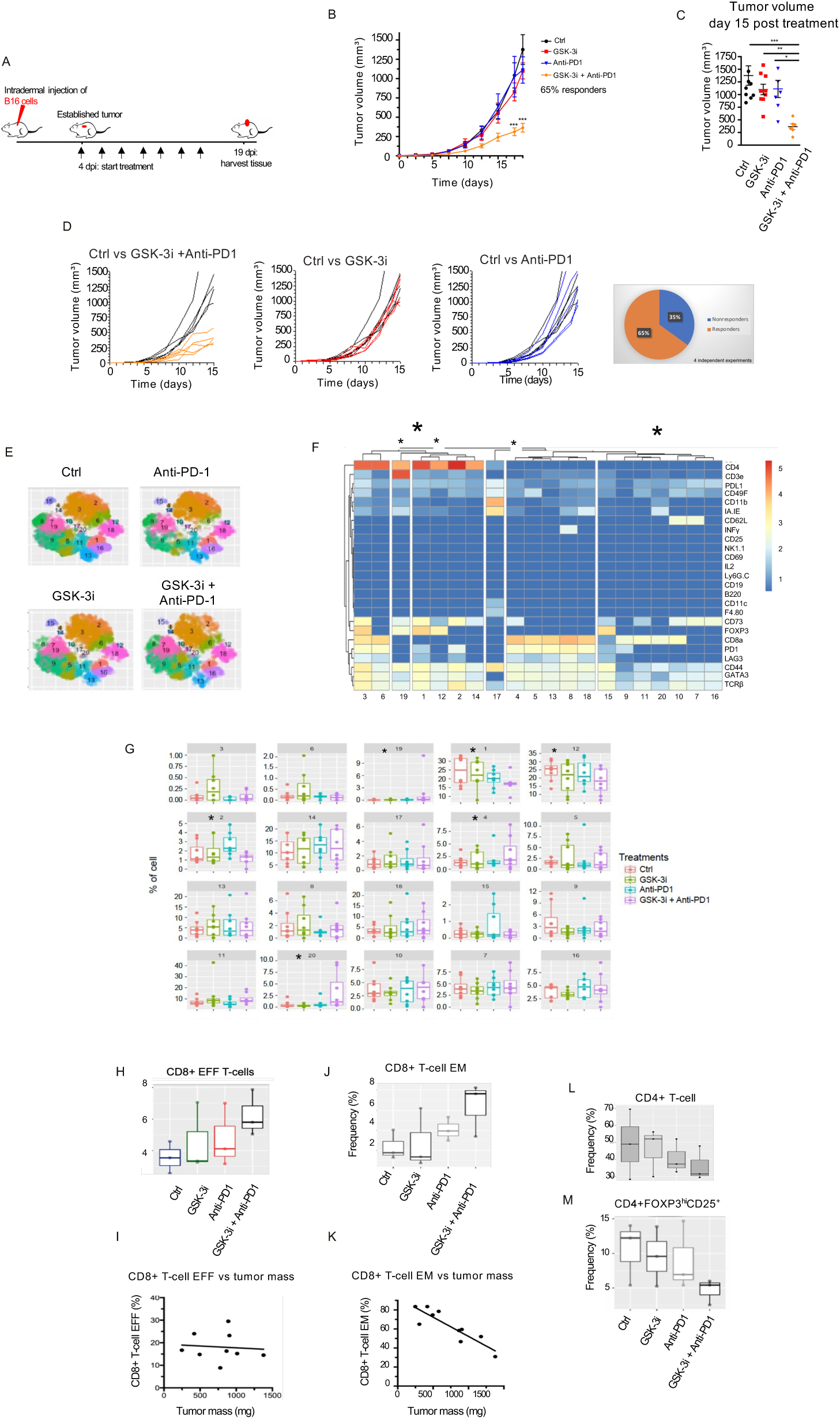
CyTOF and flow cytometric analysis of subsets of TILs induced by GSK-3 SMI enabling of anti-PD-1 therapy to resist tumor growth. (**A**) Schematic of experimental design. (**B,C**) Tumor volume quantification at day 16 post-implementation under indicated conditions (n=5-10, data pooled from ≥2 independent experiments). GSK-3i licenses anti-PD-1 to control the growth of the B16 F10 R1 tumor, otherwise resistant to monotherapy. (**D**) Left panels: Representative spider plots showing tumor progression; Right panel: Circular plots depicting proportions of responders versus non-responders following B16F10 implantation and anti-PD1 treatment with (orange) or without GSK-3 SMI (blue) (data pooled from 4 independent experiments). (**E**) Phenograph visualization of tumor-infiltrating immune cell populations by CyTOF analysis (n=5-10). (**F**) Representative heatmap showing key subsets of TILs altered by mono and combination therapy. (**G**) boxplots showing distribution of immune cell subsets identified by CyTOF. (**H**) Quantification of the percent frequency of CD8+ effector (EFF) TILs and (**I**) their lack of an effect on tumor mass. (**J**) Quantification of the percent frequency of CD8+ effector memory (EM) T cells among TILs and (**K**) the correlation analysis between CD8+ TEM TIL frequency and tumor mass (n=3). (**L)** Quantification of total CD4+ T cells and (**M)** regulatory T cells (CD25+FOXP3+CD25+), respectively (n=3).

To define TILs, we used Time-of-Flight (CyTOF) mass cytometry with a panel of 15 monoclonal antibodies (**Fig. 3E-L**). From this, 20 distinct cellular subsets within the anti-TCRβ population became apparent as shown with phenograph illustrations (**Fig. 3E**), heat maps (**Fig. 3F**) and boxplot representations (**Fig. 3G**). Despite this, although there were various upward and downward trends, only two clusters achieved statistical significance (<0.05) (**Supplementary Figure S4**). Cluster 20 showed an increase which corresponded to a CD8+ subset with the phenotype CD44+IA-IE+CD11b-CD73-CD62L- and PDL1+ (p=0.013). This effector CD8+ subset was negative for PD-1 and LAG-3 suggesting an absence of transitory or terminal exhaustion (**Fig. 3F**). By contrast, Cluster 1 in the CD4+ subset showed a decrease with the CD4+FoxP3+CD25+CD73+ suppressor phenotype (CD4+CD44+ CD25+, FoxP3+, CD73+, PD-1+LAG3+ IA-IE+CD49F+PDL1+) (p=0.044).

Using a less stringent standard, there was a trend toward a decrease in the presence of CD4 cells in Cluster 19, with a p-value of 0.09. Cluster 19 shared the expression of markers with the Treg Cluster 1 but lacked expression of PD-1 and LAG3. FoxP3+ Cluster 12 was similar to Cluster 19 but lacked expression of CD49F (integrin alpha 6) and had lower levels of CD73 inhibitory receptor CD73 (p value=0.127). Furthermore, CD8+ Cluster 4 showed an increase with a p value of 0.149. It was like Cluster 20 but expressed exhaustion markers PD-1, LAG3, and CD73. CD73 defines a highly suppressive Treg population (74, 75). Overall changes were seen in an increase in CD8+ TILs and a decrease in various regulatory T cell (Treg) suppressor due to combination therapy.

We also employed standard flow cytometry to complement the CyTOF readings (**Figs. 3H-L**). This combined approached confirmed an increase in CD8 effector TILs (**Fig. 3H**) effector-memory (EM) CD8^+^ T-cells with combination therapy (**Fig. 3J**). Interestingly, increased frequency of CD8+ EM TILs was correlated with a reduced tumor size (**Figs. 3K**). Surprisingly by contrast, there was no correlation between CD8+ EFF TILs and tumor regression (**Fig. 3I**). Analysis by both CyTOF and conventional flow cytometry revealed a decreasing trend in the proportion of CD4+ tumor-infiltrating lymphocytes (TILs) **(Fig. 3L),** as well as a more pronounced reduction in the CD4+FOXP3^hi^ CD25+ regulatory T cell (Treg) population with suppressive functions **(Fig. 3M).** Flow cytometry analysis using fluorescent antibody staining corroborated the reduction in PD-1+ cells within the Treg population and further revealed a decrease in the Treg population expressing TIM-3, for which metal-conjugated antibodies were unavailable (**Supplementary Figure S5**).

### GSK-3 licenses anti-PD-1 blockade to induce perforin and 7/9 granzymes for enhanced CD8+ CTL TILs

We next proceeded to evaluate gene expression patterns induced by anti-PD-1 and GSK-3 signaling through transcriptome profiling (**Figs. 4A-C**). Using a threshold of at least 2-fold higher expression with p<0.01, a Venn diagram illustrated the genes differentially expressed under GSK-3 inhibitor monotherapy, anti-PD-1 monotherapy, and combination therapy (**Fig. 4A**). We assessed control vs anti-PD1 monotherapy, control vs GSK-3i monotherapy, and control vs combination therapy. The combination therapy group was further divided into genes from tumors of highly responsive (i.e., termed responsive) and more poorly responsive (i.e., termed non-responsive), where responsive showed a greater than 50% reduction in tumor size relative to the mean for untreated mice. Volcano plots showed an increase and decrease in expression of different genes (x-axis shows the change magnitude while the y-axis shows statistical significance). Anti-PD-1 and GSK-3i induced a unique set of genes (162 vs 109, respectively), as well as an overlapping 3 genes. Combination therapy showed that the inhibition of GSK-3 allowed anti-PD-1 immunotherapy to express additional genes. Within this group, combination therapy induced 910 unique genes in the responsive mice and 176 unique genes in the non-responsive mice. This approach demonstrated that GSK-3 inhibition licenses anti-PD-1 blockade to induce a unique set of altered genes in immunotherapy. Further, the gene regulatory network showed an enrichment of genes associated with immune activation and inflammatory response pathways. (**Supplementary Figure S6**).

**Fig. 4:**
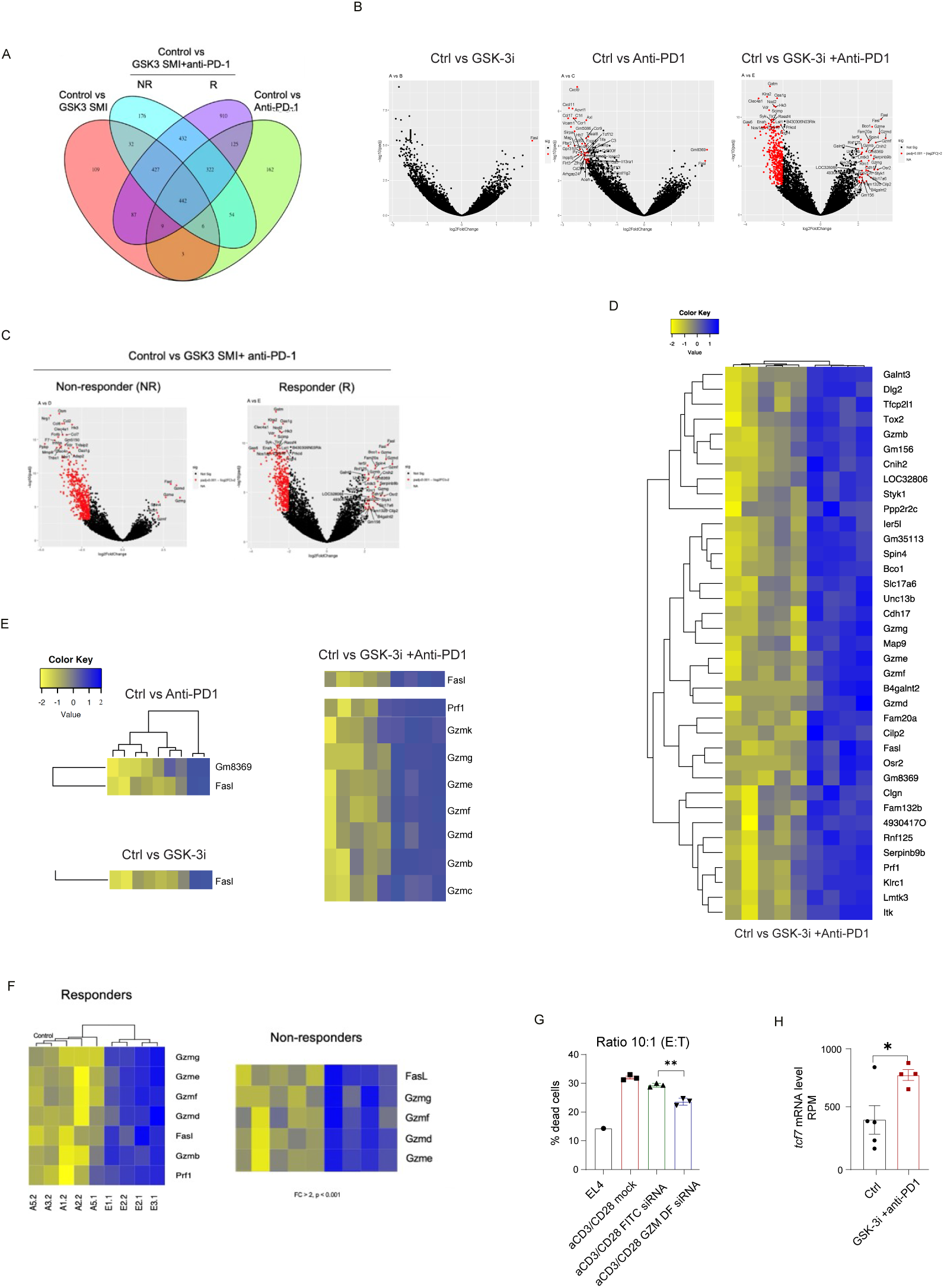
Transcriptome profiling of gene expression in TILs in response to GSK-3-anti-PD1 cooperation in cancer therapy. Analysis of differentially expressed genes in CD8+ tumor-infiltrating lymphocytes (TILs). (**A**) Venn diagram of differentially expressed genes in CD8+ TILs after RNA sequencing. (**B-C**) Representative volcano plots from RNA sequencing. Genes with a log2-fold change in expression greater highlighted in red. (**D**) Representative heatmap of overexpressed genes with a log2-fold change in expression greater than 1 and a p-value of 0.001. (**E-F**) Representative heatmap of cytotoxic genes with a log2-fold change in expression greater than 1. (**G**) Quantification of the proportion of dead cells following a killing assay with GZMD and F siRNA knockdown. (**H**) Quantification of *tcf7* mRNA levels (reads per million, RPM) in CD8+ TILs. Data are shown as mean ± SEM. Statistical significance was determined by unpaired t-test (*p < 0.05, **p < 0.01, ***p < 0.001, ****p < 0.0001, NS: not significant). n = 3 with 4-5 mice per group in each experiment.

Due to the extensive gene list, we implemented a more stringent filter requiring a minimum two-fold expression change and a p < 0.001 significance level (**Fig. 4B**). Under these conditions, volcano plot analysis revealed limited positive gene induction by GSK-3 SMI inhibition alone involving FasL (**Fig. 4B**). In contrast, while anti-PD-1 monotherapy negatively impacted the expression of numerous genes such as VCam1, CXCL9, CXCL11, and others, it positively affected only two genes, FasL and Gm8369 (upper panel). In contrast, the combination of anti-PD-1 and GSK-3 SMI synergistically induced many more genes, both negatively and positively (lower panel). Differences were also seen in TILs from responsive and non-responsive mice (**Fig. 4C**).

With a focus on the upregulated 37 genes, we noted the induction of important genes such as interleukin 2 inducible kinase (ITK) and the transcription factor Tox2. ITK regulates cytokine production and T-cell differentiation (76), while Tox2 is required for metabolic adaptation and tissue residency of immune cells (77). However, in addition was the remarkable induction of perforin and 7/9 granzymes from within the mouse genome (**Fig. 4D**). These included granzymes, GZMK, GZMG, GZME, GZMF, GZMD, GZMB and GZMC needed for optimal CD8+ CTL killing function (**Fig. 4E**). Perforin is a pore forming cytolytic protein found in the granules of CTLs, while granzymes enter via perforin to induce cell death (78, 79). This effect is, to our knowledge, unprecedented.

In comparing the highly responsive (i.e., termed responsive) and poorly responsive (i.e., termed non-responsive) mice, the TILs from the poorly responding mice showed an increase in the expression of GZMG, GZME, GZMF, and GZMD, but no significant increase in GZMK, GZMB, or perforin expression (**Fig. 4F**). A trend to increase was observed but it was not statistically significant. Perforin, a critical pore-forming protein essential for CTL-mediated cytotoxicity, resides within cytotoxic granules and is influenced by PD-1 internalization (25). Furthermore, GZMB has been previously demonstrated to play a pivotal role in CD8+ T cell effector function of TILs (80).

To confirm the role of the lesser-known granzymes in CD8+ T-cell-mediated killing, siRNA to down-regulate GZM D and F was assessed in an OT-1 CTL killing assay (**Fig. 4G**). OT-1 T-cells were activated with anti-CD3/CD28 for 24 hours and electroporated with siRNAs targeting both GZMD and GZMF. Cytotoxic CD8+ T cells were cultured for 48 hours before being used in a killing assay against EL4 target cells. While a control siRNA with a fluorescent label (FITC-scrambled) had minimal impact on killing activity, siRNAs targeting GZMD and GZMF reduced killing by more than 20% compared to the FITC-scrambled control and cells without RNA transfection (mock). These findings showed roles for GZMD and GZMF in the killing efficiency of cytotoxic CD8+ T cells. Furthermore, building upon the previous analysis of TCF-1+ stem-like T cells, the combined treatment of anti-PD-1 and GSK-3 SMI significantly increased *tcf7* expression (**Fig. 4H**).

### GSK-3KD licenses anti-PD-1 induction of CD8 multi-granzyme expression

These findings were further corroborated by comparing the response of WT vs GSK-3 KD mice to anti-PD1 blockade (**Fig. 5**). While anti-PD1 failed to significantly inhibit B16 F10 R1 tumor growth in WT mice, it reduced tumor growth by more than 75% in dLCK-CRE mice. This was evident in both tumor volume (**Fig. 5A**) and by the measurement of tumor weight at day 16 post-implantation (**Fig. 5B**). Further, flow cytometry analysis of a representative experiment showed that anti-PD1 increased the frequency of CD8+ TILs from 37% to 46.3% in WT mice (**Fig. 5C**). The increase in CD8+ TILs was further augmented to 60.2% in GSK-3 KD mice. Consistent with the increase in CD8+ T-cells, the total number of CD8+ TILs increased in WT in response to anti-PD1 from 2.4 x 10^6^ cells/g. However, in the KD mice, this increased to 9.2 x 10^6^ cells/g (**Fig. 5D**).

**Fig. 5:**
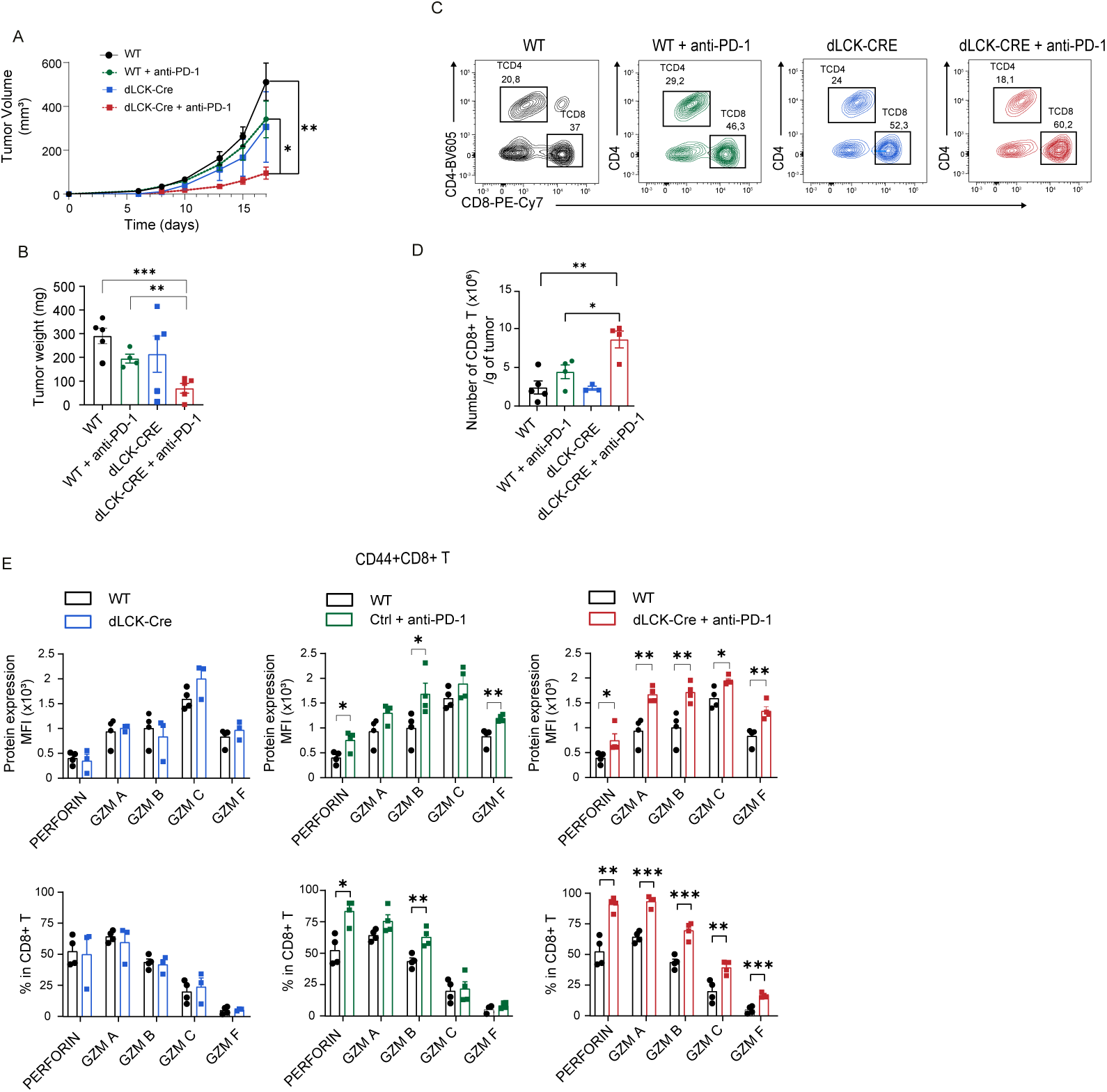
GSK-3 KD licences anti-PD-1 blockade to generate super-armed CD8+ CTL TILs in tumors. B16F10 implanted WT and dLCK-CRE mice were injected or not with anti-PD1. (**A**) Tumor volume in control (ctrl) and dLck-Cre mice with or without anti-PD-1 treatment. (**B**) Tumor weight 16 days post-implantation. (**C**) Representative flow cytometry plots of CD4 and CD8+ TILs following checkpoint blockade in WT and dLck Cre mice. (**D**) Numbers of CD8+ TILs per gram of tumor tissue. **(E**) Expression and proportion of perforin, granzyme A, B, C, and F in CD44+CD8+ TILs in WT and dLck Cre mice. Data are presented as mean ± SEM. Statistical significance was determined by unpaired t-test (*p < 0.05, **p < 0.01, ***p < 0.001, ****p < 0.0001, NS: not significant). n = 3 experiments with 4--5 mice per group.

FACS staining of granzymes then showed that anti-PD-1 in GSK-3 KD mice significantly upregulated the expression of perforin and multiple granzymes, GZMA, GZMB, GZMC, and GZMF (as assessed by available anti-granzyme antibodies) (**Fig. 5E**). Other granzymes did not have suitable antibodies which worked in flow cytometry. Nevertheless, while anti-PD1 monotherapy resulted in an increased frequency of GZMB-expressing CD8+ T-cells in WT mice, combination of anti-PD-1 in GSK-3 KD mice significantly upregulated the expression of perforin, GZMA, GZMB, GZMC, and GZMF in terms of mean fluorescence intensity (MFI) of expression (upper panel) and the percentage of CD8+ T-cells expressing each granzyme (lower panel). These data demonstrate that GSK-3 inhibition synergizes with anti-PD1 therapy to enhance the cytotoxic capacity of CD8+ T-cells by upregulating the expression of a broad spectrum of granzymes, thereby facilitating more effective tumor eradication.

### GSK-3 alters CD4+ Treg vs T-help in tumors and regulates CD8+ CTL dependency on CD4+ T cell help

Given that CD8+ T-cell differentiation can be facilitated by CD4+ T-cell help (4-6), a key next question was whether the induction of GZMs was due to a role of GSK-3 in regulating CD4-CD8 cooperation where help is provided to CD8+ T-cells (**Fig. 6).** The downregulation of GSK-3 did not affect the overall numbers of CD4+ TILs (**Fig. 6A**). It did however increase the numbers of T-helper cells relative to CD4+ FoxP3+ suppressor cells in the TIL population **(Fig. 6B**). In a representative FACs plot, anti-PD-1 therapy induced an increase in the presence of CD4+ FoxP3+ regulatory TILs in WT mice from 36.8 to 59.8% of CD4+ CD44+ TILs, as reported (81-84). At the same time, the percentage of FoxP3 negative CD4+ TILs decreased. In striking contrast, in dLck-CRE mice, anti-PD-1 cooperated to increase the percentage of T-helper cells from 40.2 in WT to 73.8, while concurrently decreasing the presence of Tregs from 59.8 in WT to 26.4%. These data, consistent with the CyTOF analysis, showed that GSK-3 plays a pivotal role in regulating the relative presence of CD4+ T helper vs Treg TILs in response to anti-PD-1 therapy.

**Fig. 6:**
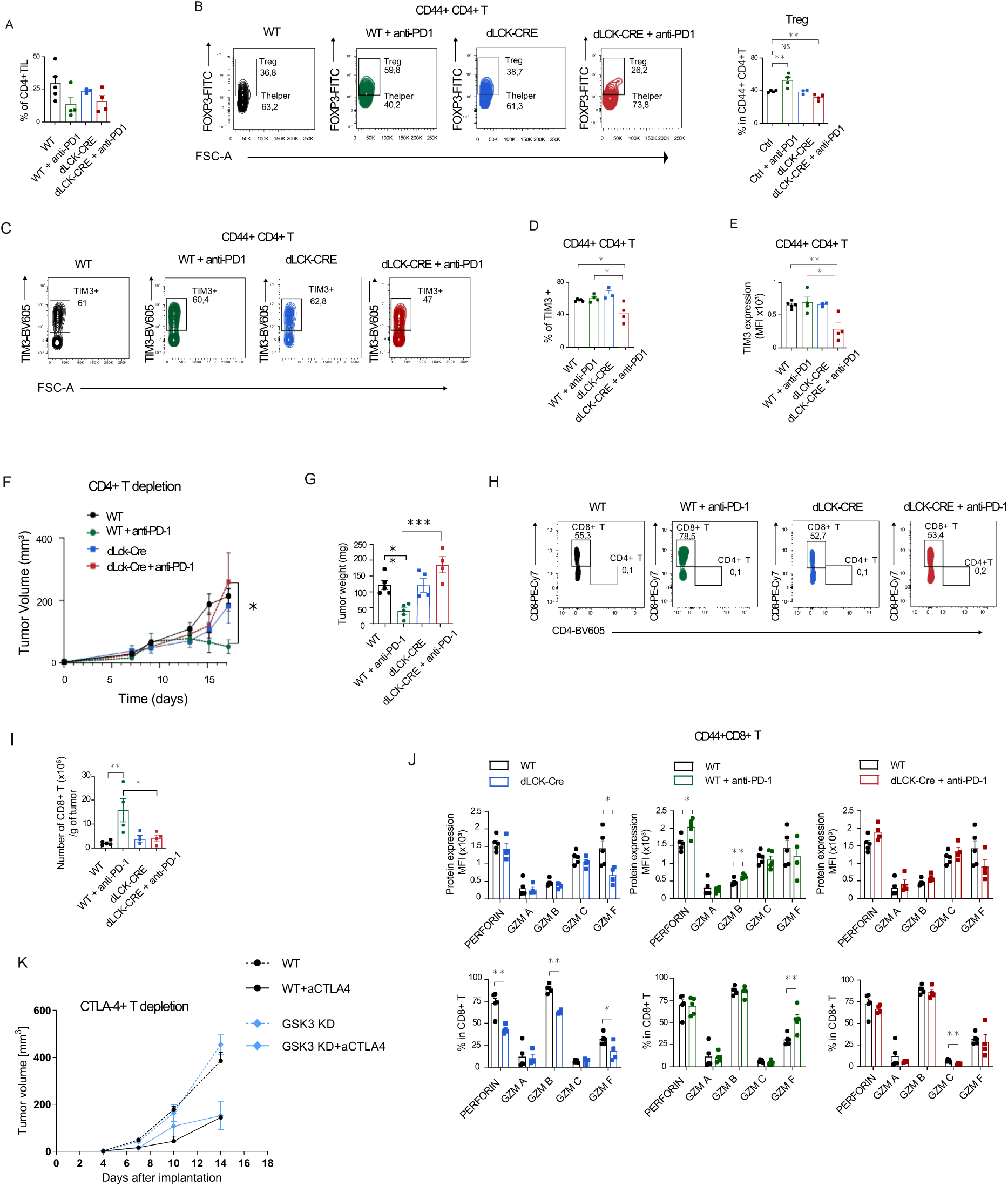
GSK-3 regulates CD4+-CD8 cooperation needed for the efficacy of PD-1 blockade to generation multi-granzyme CTLs in tumors. (**A**) Quantification of CD4+ tumor-infiltrating lymphocytes (TILs) in wild-type (WT) and dLCK-Cre mice with or without anti-PD-1 treatment. (**B**) Representative flow cytometry analysis of CD4+ Treg and helper TILs in WT and dLck-Cre mice following anti-PD-1 therapy. (**C-E**) Representative flow cytometry plots (**C**) and histogram quantification **(D, E**) of TIM3+ expression on CD44+CD4+ TILs with or without anti-PD-1 treatment. (**F, G**) Measurement of tumor volume (**F**) and weight (**G**) following CD4+ T cell depletion. (**H**) Representative flow cytometry plots of the presence of CD8+ TILs after CD4+ T cell depletion. (**I**) Measure of the numbers of CD8+ TILs/g of tumor after CD4+ T cell depletion. (**J**) Quantification of the loss of the induced expression of perforin, granzyme A, B, C, and F expression in CD44+CD8+ TILs after CD4+ T cell depletion in GSK-3 KD mice, shown as mean fluorescence intensity (MFI) (upper panel) and percentage (lower panel). Quantification of target cell killing with or without anti-PD-1 treatment. (**K**) Depletion of Tregs with anti-CTLA-4 reversed the inhibition of tumor regression in WT mice while not affecting the ability of dLck-Cre mice to limit tumor growth. Data are presented as mean ± SEM. Statistical significance was determined by unpaired t-test (*p < 0.05, **p < 0.01, ***p < 0.001, ****p < 0.0001, NS: not significant). n = 4-5 mice per group.

Further, anti-PD-1 treatment in dLCK-CRE mice also led to a significant reduction in both the frequency and expression of the inhibitory receptor TIM3 on CD44+CD4+ TILs, as demonstrated by flow cytometry analysis. This decrease in expression was evident in the FACS plots (**Fig. 6C**) as reflected in a decrease in the percentage of TIM3+ cells (**Fig. 6D**), and level of expression (i.e., MFI) (**Fig. 6E**). Tim-3 expression on CD4+ T-cells promotes T-cell exhaustion, while expression of Treg cells correlates with greater suppressive ability (85).

Given the changes in the composition of the CD4 subset, we next depleted CD4 cells by injecting mice at various time points to remove CD4+ T-cells before and during anti-PD-1 therapy. Interestingly, CD4 depletion allowed anti-PD-1 therapy to reject the B16-F10 R1 tumors in wild-type (WT) mice (**Fig. 6F**). This was evident from the reduced tumor volume (**Fig. 6F**) and tumor weight (**Fig. 6G**). Since anti-PD-1 therapy alone was normally unable to promote tumor rejection and typically increased the presence of regulatory T cells (Tregs), these observations suggested that the loss of CD4+ Tregs through anti-CD4 depletion enabled anti-PD-1 therapy to reject tumors, which was not observed when CD4+ T-cells were present.

Conversely, in dLCK-CRE mice, with greater numbers of CD4 T-helpers, the depletion of CD4 TILs caused a complete loss in the ability of anti-PD-1 to promote tumor rejection (**Fig. 6F, G**). This was seen both in tumor volume (**Fig. 6F**) and tumor weight (**Fig. 6G**). In this scenario, anti-PD-1 lost its ability to promote an increase in the presence of CD8+ TILs as shown in a representative FACs plots (**Fig. 6H**) or in terms of numbers of CD8+ T-cells/g of tumor (**Fig. 6I**). In addition, the loss of CD4 help in the GSK-3 KD mice resulted in a complete loss in the increase in the expression of the multiple granzymes in the CD8+ TIL population (**Fig. 6J**). This loss was seen in terms of protein expression (i.e., MFI) (upper panel) and in the percentage expression (lower panel) for perforin, GZMA, GZMB, GZMC and GZMF.

Confirmation that the suppression of the anti-tumor response due to the presence of Tregs in the TIL population from WT mice was seen by the ability of anti-CTLA-4 depletion to also allow a reduction in tumor size (**Fig. 6J**). Anti-CTLA-4 is effective in depleting CD4+ Tregs (86). By contrast, anti-CTLA-4 had no effect on the regression of the B16 F10 R1 tumor in the GSK-3KD mice. Since anti-CD4 did reverse the enhanced regression in tumors in the dlck-CRE mice (**Fig. 6F**), these data indicate that the effect of anti-CD4 on tumor regression in T-cells with reduced expression of GSK-3 was due to the loss of CD4+ T-helper cells.

To test for a potential difference in dendritic cells (DCs), *in vitro* derived bone marrow derived DCs from dLCK-CRE vs WT mice supplemented with granulocyte-macrophage colony-stimulating factor (GM-CSF) were found to exhibit comparable surface antigen expression profiles (**Supplementary Figure S7**). Overall, these results demonstrate that GSK-3 expression plays a pivotal role in determining the relative presence of T-helper cells versus Tregs in tumors in response to anti-PD-1 treatment and consistent with this, the dependency of CD8+ T-cells on CD4+ T-cell help for their presence in tumors and their expression of multiple granzymes needed for tumor killing.

## Discussion

Despite its importance in T-cell signaling, the full role of GSK-3 in immune checkpoint blockade has yet to be explored. In this study, we show that GSK-3 normally limits the ability of anti-PD-1 to elicit the full development of CD8 CTL responses against tumors. Further, this involves the regulation of the balance between the presence of regulatory and helper T-cells in their regulation of CD8 expression of cytolytic granzymes needed to kill tumors. The genetic knockdown of GSK-3 increased memory TCF-1+ T-cells with reduced overall exhaustion for more potent anti-viral and tumor immunity. GSK-3 knockdown and inhibition licensed anti-PD-1 to overcome tumor resistance in a process that down-regulated CD4+ Tregs while increasing the presence of effector-memory CD8+ T-cells. This synergy resulted in the generation of super-armed killer T-cells expressing an unprecedented 7/9 cytolytic granzymes (GZMs K, G, E, F, D, B, C). Importantly, reduced GSK-3 expression was essential for CD4+ helper collaboration with CD8+ T-cells to express these unprecedented numbers of cytolytic granzymes. Our findings offer a novel therapeutic strategy for cancer immunotherapy, with the potential to improve treatment outcomes by optimizing CD4-CD8 T-cell crosstalk needed to increase GZM expression.

Our initial findings showed that GSK-3 KD favored the generation of TCF1+ progenitor memory cells. This feature was first seen in LCMV Cl13 infection, showing a continued increase in both central and effector memory T-cells as well as a preferential increase in the presence of MPECs relative to SLECs. MPECs play a key role in developing immunological memory to respond to future infections (87, 88). This increase in TCF1+ progenitors in GSK-3 inhibited conditions concurrently led to a reduction in T-cell exhaustion using in vivo and in vitro assays. Small chemical inhibitors have previously used (89), although there have been no reports on the connection between GSK-3 and memory progenitors using genetically altered CD4+ and CD8+ T-cells. A selective KO in CD8+ T-cells showed reduced exhaustion but failed to establish a link to the generation of TCF-1+ progenitor T-cells (90). Surprisingly, the same subset showed an increase in the expression of the cell cycle marker Ki67 in GSK-3 KD but not wild-type mice. Further, in our study MPECs showed an increase in the expression of GLUT-1 and an increase in glucose uptake. These observations place the GSK-3 kinase upstream in a signaling pathway in the control of the generation of TCF-1+ progenitor memory T-cells (91, 92).

Since the presence of progenitor TCF-1+ T-cells is required for successful anti-PD-1 therapy (26-28), we next tested whether GSK-3 inhibition with small molecule inhibitors (SMIs) or its genetic downregulation could cooperate with anti-PD-1 in eliminating a tumor that is normally resistant to checkpoint blockade. In the B16 F10 R1 model, tumor growth was unaffected by monotherapy but was reduced by more than 70% when assessed on day 16 with the combination treatment. Furthermore, tumor regression was associated with a significant shift in the tumor microenvironment. Specifically, CyTOF analysis showed a loss of highly suppressive FOXP3hi CD25+ Tregs concurrent with an increase in CD8+ effector (EFF) and effector memory (EM) CD8+ TILs. The increase in EM cells in the tumor model was correlated with a reduction in tumor mass. The lack of correlation with EFF CD8+ T-cells might relate to their more terminal differentiation and shorter lifespan needed to sustain a response to tumor neoantigens. GSK-3 expression plays a central role in determining the relative presence of Tregs and CD8 effector memory cells in tumors in response to anti-PD-1 therapy.

Transcriptomics also showed that GSK-3 down-regulation controlled the response of the T-cells to anti-PD-1 by licensing blockade to regulate many more genes that normally induced by anti-PA-1 alone. The effect was also selective as shown by a skewed effect in preferentially affecting immune associated genes. Further, and of special interested was selective effect on the unprecedented increase in the expression of granzymes needed to kill tumors. Of the 37 genes induced by combination therapy, these included perforin, a pore-forming protein crucial for CTL cytotoxicity (93), and 7/9 granzymes in the mouse genome. Therefore, at least 8 of 37 induced genes were genes that controlled CTL killing. This included the well-known GZMB as well as less well-studied granzymes GZMK, GZMG, GZME, GZMF, GZMD, and GZMC. Notably, cells expressing high levels of GZMF are distinct from those expressing GZMA and GZMB in their mechanism of eliminating cancer cells (31). GZMC and GZMK employ a different caspase-independent pathway for killing (32). This upregulation of multiple granzymes is unprecedented in the literature and indicative of a ‘super-arming’ of cytolytic T cells, enhancing their capacity for tumor destruction.

To achieve this remarkable effect, we also uncovered the cellular mechanism by showing that levels of GSK-3 expression determine the relative presence of T-helper cells versus Tregs in tumors in response to anti-PD-1 treatment. Further, it determined the dependency of CD8+ T-cells on CD4+ T-cell help for their presence in tumors for the expression of the multiple granzymes needed for tumor killing. We observed an increase in the presence of Tregs seen in WT mice in response to anti-PD-1 which failed to occur in GSK-3KD mice. Remarkably instead, the reduced level of GSK-3 allowed anti-PD-1 to support the increased presence of CD4+FoxP3 negative helper TILs. In turn, anti-CD4 depletion experiments showed that this CD4+ T cell help was needed for the regression of tumors, the increased presence of CD8+ TILs and crucially, for inducing the multiple GZMs including GZMK, GZMG, GZME, GZMF, GZMD, and GZMC machinery of GSK-3 KD CD8+ CTLs. This suggests that GSK-3 is a key player in the development of poorly performing CD8 T-cells in the absence of CD4 T-cell help (4, 5, 7)..

The depletion of Tregs with anti-CTLA-4, in turn, mirrored the effect of anti-CD4 in allowing WT mice to clear the tumor. This showed that the anti-CD4 depletion effects were most likely related to its depletion of Tregs in WT mice, which normally suppresses tumor clearance. However, in the case of the dLckCre mice with reduced levels of GSK-3 and the reduced presence of Tregs, anti-CTLA-4 had no effect on the enhanced ability of the mice to regress tumors. This supports the notion that anti-CD4 reversed tumor size reduction in dLckCre mice by eliminating CD4+ helper T-cells. This pathway by which GSK-3 controls anti-PD-1 effects on CD4-CD8 cooperation for the relative presence of suppression versus help and the development of super-armed CD8+ TILs is a new pathway in the regulation of tumor immunotherapy.

One contributing factor not yet reported is that the reduction in GSK-3 caused T-cells to be more reliant on glycolysis relative to oxidative phosphorylation for their energy requirements. The effect on extracellular acidification rate (ECAR) was more striking than on the oxygen consumption rate (OCR) with a lower OCR/ECAR ratio. This reliance on glucose uptake was also evident with the greater glucose uptake and GLUT-1 expression on GSK-3 KD cells, where this effect was most evident in GP33+ CD8+ MPECs. This observation, combined with the greater cell division evident in MPECs, as seen with Ki67 expression (data not showed), is consistent with the shift in T-cells from the primarily use of oxidative phosphorylation to glycolysis upon activation (94, 95). In addition, stem-like progenitor T-cells primarily utilize glycolysis to generate ATP (96). Although not reported before for T-cells, B-cell responses to the co-stimulatory receptor CD40 also engage glycolysis (97). In another study, IL-24 has been reported to inhibit GSK3β is mediated through PKA activation, activating glycogen synthase and decreasing intracellular glucose levels (98).

Overall, our findings in this paper related to GSK-3 regulation of CD4-CD8 collaboration in the generation of CD8 CTLs could provide a mechanism to explain the past findings showing that GSK-3 can promote CAR function and tumor rejection (42, 54-56, 59-61). The one exception was the generation of mice using a granzyme B (GZMB)-Cre for GSK-3 deletion that showed an increase in tumor growth (90). However, this difference may relate to the fact that the GZMB-Cre system also targets NK cells and cytolytic CD4+ T cells, regulatory suppressor CD4+ T cells (Tregs), Group 1 ILCs, mast cells, and macrophages (99). Overall, our findings underscore the importance of the serine/threonine kinase GSK-3 in governing the nature of the immune response in checkpoint blockade, both in terms of its control of the cooperation between CD4 and CD8 T-cells and enhanced GZM expression in anti-PD-1 therapy.

## Acknowledgements

We thank Dale long and his team from NIH Tetramer Core Facility at Emory for providing GP33 tetramers and Fredéric Duval and Anne-Mari Aubin from the Flow Cytometry Core Facility of CR-HMR for their help. This project was supported by the Canadian Institutes of Health Research Foundation grant (159912) (PI) and National Institutes of Health grant RO1 AI049466 (co-PI). CER holds a majority stake in ImmunoAb Res Inc (Canada).

## Methods

### Ethic approval

Animal experiments were approved by the Animal Care Use and Review Committee of Center Research -Hospital Maisonneuve Rosemont (CR-HMR) (ethical approval no 2021-2516, 2023-3032 and 2024-3547) from University of Montreal. For mouse tumor experiments, all tumor sizes/burdens were permitted by the CR-HMR of Montreal University. Maximal tumor burden permitted by the CR-HMR (1.5 cm3) was not exceeded in this study.

### Key resources

A complete list of the reagents and antibodies used in this manuscript can be found in supplementary table S1.

### Animals

#### Generation and Utilization of GSK-3αflox/flox βflox/flox mice

Two generations of GSK-3αflox/flox βflox/flox mice were produced: one in Montreal and another in our laboratory at the University of Cambridge, UK. The mice generated in Montreal were used exclusively for most experimental procedures. Splenocytes from the University of Cambridge were transported to the University of Leeds, courtesy of Dr. Alison Taylor, who kindly arranged their overnight shipment on ice to Montreal. These splenocytes were utilized in Supplementary Figure 3. GSK-3αflox/flox and βflox/flox mice were crossed to generate double-negative mice in Montreal. For both the Montreal and Cambridge cohorts, GSK-3αflox/flox βflox/flox mice were bred with distal dLCK-Cre mice (B6.Cg-Tg(Lck-icre)3779Nik/J, Jackson Laboratory, Maine) to delete both GSK-3α and GSK-3β isoforms. However, in the case of the Montreal mice, the deletion was incomplete, resulting in the creation of GSK-3 knock-down mice. All mice were maintained in an animal facility with a 12-hour light-dark cycle and had unrestricted access to food and water throughout the study. The study adhered to ethical guidelines for animal research. Sex was not a factor in the study design or analysis, as the study was not intended to investigate sex differences.

#### Viral infections

LCMV Armstrong and Clone 13 strains were propagated in BHK-21 cells and their titers determined using a plaque assay on Vero cells (100). Mice were infected with either LCMV Armstrong via intraperitoneal (i.p.) injection at a dose of 2 × 10^!^ plaque-forming units (PFU) or with LCMV Clone 13 via intravenous (i.v.) injection at the same dose. Following infection, mice were sacrificed at either day 7 (for LCMV Armstrong) or day 30 (for LCMV Clone 13) to assess the effects of infection.

#### Cell lines

B16F10 tumor cells were maintained in DMEM supplemented with 10% FCS, 1% L-glut and 1% Pen/Strep. Tumor cells (2x10^5^) cultured for less than two weeks were implanted subcutaneously in the flank of recipient mice using 29G1/2 syringes. Tumor size was monitored using a manual caliper and tumors were excised before exceeding the volume permitted by CR-HMR ethic committee.

#### Spleen Processing and T Cell Activation

Spleens were harvested from 8-to 16-week-old mice and mechanically dissociated in a 6-well plate containing 2 mL of red blood cell lysis buffer (ACK buffer). An additional 2 mL of lysis buffer was added, followed by a 5-minute incubation at room temperature. The reaction was quenched with 20 mL of phosphate-buffered saline (PBS), and the suspension was filtered through a 70 μm cell strainer. Cells were pelleted by centrifugation at 350 × *g* for 5 minutes.

Splenocyte pellets were resuspended in 1 mL of FACS buffer (PBS supplemented with 2 mM EDTA and 2% fetal bovine serum [FBS]). Viable cells were quantified by Trypan blue exclusion using a Neubauer hemocytometer. CD3+ T cells were isolated via magnetic bead-based negative selection (StemCell Technologies). Following isolation, 2 × 10⁶ T cells/mL were seeded in 1 mL of activation medium (RPMI-1640 with L-glutamine, 10% FBS, 1 M HEPES, and 0.1% β-mercaptoethanol) in a 48-well plate. Cells were activated with 1 μg/mL anti-CD3ε antibody (clone 145-2C11) for 24 hours at 37°C under standard culture conditions.

#### Antibody Administration Protocol

Anti-PD-1 antibodies (Leinco) and SB415286 (Selleckchem) were reconstituted at 1 mg/mL in sterile phosphate-buffered saline. Mice were administered 200 μg of the antibody solution via intraperitoneal injection every 48 hours, starting on day 6 post-tumor implantation and continuing until tumor resection on day 16<u>^[1]^</u>.

#### CD4+ and CTLA-4 T-cell Depletion

To deplete CD4+ T cells, mice received 300μg of anti-CD4 antibody (Leinco) via intraperitoneal injection 24 hours prior to tumor implantation (day -1), with subsequent doses administered on days 7 and 14 post-implantation. This regimen ensured sustained depletion throughout the experimental timeline. The small regime was used in the depletion of Tregs with anti-CTLA-4. C57BL/6 mice were injected with 1 × 10^5^ B16F10 melanoma cells. In the αCTLA4 groups, mice received 200 μg of CTLA4 (clone 9d9) blocking antibody (Leinco) IP at 4, 7, and 10 days thereafter. Tumors were measured by caliper every three days, and tumor volume was calculated as length x width x width/2. At the time of sacrifice for analysis, mice were euthanized using CO2 and subsequent cervical dislocation.

#### Glucose Uptake and Metabolic Profiling in Activated T Cells

Splenocytes were isolated and activated with 1 μg/mL anti-CD3ε antibody (clone 145-2C11) for 48 hours at 37°C under standard culture conditions. For glucose uptake assessment, activated cells were incubated with 2-NBDG (Thermo Fisher Scientific, #N13195) at a 1:1,000 dilution in phosphate-buffered saline (PBS) for 1 hour at 37°C, followed by fluorescence quantification via flow cytometry.

#### Seahorse XF24 Metabolic Analysis

Oxygen consumption rate (OCR) and extracellular acidification rate (ECAR) were measured using an Agilent Seahorse XF24 Extracellular Flux Analyzer. T cells were isolated, activated as above, and seeded onto Cell-Tak-coated Seahorse cell culture plates. Prior to analysis, cells were equilibrated for 90 minutes in Seahorse assay medium supplemented with 10 mM glucose and 1 mM sodium pyruvate<u>^[3][4]^</u>.

Mitochondrial respiration was evaluated using the Mito Stress Test Kit with sequential injection of 0.5 μM rotenone/antimycin A (Rot/AA) and 2-NBDG (50 μM). Glycolytic capacity was assessed via the Glycolytic Rate Assay Kit using 1.5 μM oligomycin, 2 μM carbonyl cyanide-4-(trifluoromethoxy)phenylhydrazone (FCCP), and 0.5 μM Rot/AA<u>^[5][4]^</u>. Data were normalized to cell density (2 × 10⁵ cells/well) and analyzed using Wave software (Agilent Technologies)<u>^[3][6]^</u>.

### Reagents and antibodies

A complete list of the reagents and antibodies used in this manuscript can be found in supplementary table S1.

#### Cell Preparation and Immunophenotyping

Organs were harvested from 8- to 16-week-old mice. Inguinal lymph nodes (LN) and thymus were mechanically dissociated through a 70 μm cell strainer using the plunger of a 5 mL syringe. The strainer was washed with 5 mL of FACS buffer (PBS supplemented with 10 mM EDTA and 2% fetal bovine serum [FBS]). Tumors were mechanically disrupted on 70 μm strainers in RPMI medium on ice. Tumor lysates were centrifuged at 350 × *g* for 5 minutes and resuspended in PBS. The tumor lysate in PBS was layered onto lymphocyte separation medium (Corning) and centrifuged at 2000 rpm for 20 minutes at room temperature to isolate tumor-infiltrating lymphocytes (TILs).

#### Cell Blocking and Staining

Isolated cells from spleen, LN, thymus, and tumor were blocked with 100 μL of PBS containing 10 mM EDTA, 5% FBS, and 2.5 ng/mL purified rat anti-mouse CD16/CD32 (BD Biosciences #553141) for 20 minutes at 4°C. Cells were then centrifuged at 350 × *g* for 5 minutes and stained with primary antibodies (as detailed in the supplementary table) for 30 minutes at 4°C. Each staining mix was supplemented with 1/1000 Fixable Viability Dye eFluor™ 780. Following staining, cells were fixed with 2% paraformaldehyde (PAF) for 15 minutes at 4°C, washed twice with PBS/10 mM EDTA + 2% FBS, and then permeabilized using the FOXP3 kit (BioLegend) for 30 minutes at 4°C. Cells were stained with antibodies targeting intracellular proteins, diluted in permeabilization buffer, for 50 minutes at room temperature. After staining, cells were washed twice with PBS/10 mM EDTA + 2% FBS.

#### Flow Cytometric Analysis

Cell suspensions were analyzed using a BD LSRFortessa flow cytometer. Mean fluorescence intensity (MFI) of specific markers was quantified on T cell subpopulations using FlowJo software (Tree Star, Ashland, USA). Results were further analyzed using FlowJo to assess immunophenotypic profiles.

### Immunoblotting

Cell lysates were prepared by resuspension of spleen-purified CD8+ T cells in 1% Triton X-100 lysis buffer, followed by a 30-min incubation on ice as described (101). Lysates were centrifuged at 14,000 rpm for 30 min at 4°C and the nuclear-free supernatants were subjected to SDS-PAGE on a 10 % polyacrylamide gel. Proteins from the gel were electroblotted onto nitrocellulose membranes at 100V for 75 min, followed by 1 h of blocking with 3% BSA in TBST at RT. The nitrocellulose membranes were then incubated in the presence of the indicated primary Abs, followed by incubation with HRP-conjugated secondary Abs. Immunoreactive protein bands were visualized using an ECL reagent and autoradiography. Whenever required, nitrocellulose membranes were stripped by incubation in stripping buffer for 20 min at RT, followed by 1 h incubation with blocking buffer (3% BSA in TBST).

### Mass cytometry (CyTOF)

Mass cytometry (CyTOF) analysis was performed at the McGill University CyTOF facility following standardized protocols. Single-cell suspensions (3×10⁶ cells) were stained with 5μM Cell-ID™ Cisplatin (Fluidigm, San Francisco, CA) for viability assessment, followed by32quenching with MaxPar® Cell Staining Buffer at a 5:1 volume ratio. Following centrifugation (300g, 5 min, 4°C), cells were resuspended in 50 μL buffer and incubated with human Fc receptor blocking solution (5 μL; BioLegend, San Diego, CA) for 10 minutes at room temperature. Surface marker staining was performed using 50 μL of pre-titrated metal conjugated antibody cocktail for 30 minutes (detailed panel composition in Reference 20). After washing, cellular DNA was labeled overnight at 4°C using 1 mL intercalation solution containing 125 nM MaxPar® Intercalator-Ir in MaxPar® Fix and Perm Buffer. Prior to data acquisition, cells were washed with MaxPar® Water, resuspended with EQ™ Four Element Calibration Beads, and analyzed on a Helios™ CyTOF II mass cytometer. A minimum of 100,000 viable single cells were acquired per sample at an event rate of 300-500 events/second. Analysis was performed on using antibodies against CD45, TCR-beta as indicated in Figure 4A.

### Mass cytometry data analysis

Mass cytometry data analysis was performed using the Cytobank platform (Cytobank Inc., Mountain View, CA) following previously established protocols94, 95. All samples underwent simultaneous normalization and analysis to minimize technical variation and account for signal drift during extended acquisition periods. A standardized high-level gating strategy was implemented across all CyTOF files, incorporating sequential gates for event length, bead exclusion, cell viability, and DNA content. For comprehensive analysis, individual sample files were concatenated into a single dataset, from which population-specific data were subsequently extracted and separated into individual files for comparative analyses. Dimensional reduction was performed using t-Distribution Stochastic Neighbor Embedding (tSNE) analysis, which generated a two-dimensional representation (tSNE1 versus tSNE2) of the high-dimensional data by preserving local phenotypic relationships between cells, enabling visualization and identification of distinct cell populations based on their marker expression profiles.

### Sample preparation for RNAseq and RNA profiling

Tumor was prepared as previously described and CD8+ TILs were isolated (Stemcell Technologies) RNA was isolated using the miRNeasy Mini Kit (Qiagen), and the mRNA was purified from 1ug of total RNA using the Dynabeads mRNA DIRECT Micro Kit (Thermo Fisher Scientific). The reverse transcription used SuperScript VILO cDNA Synthesis kit (Thermo Fisher Scientific). The whole transcriptome libraries were prepared using the Ion Total RNA-Seq Kit version 2 Kit (Thermo Fisher Scientific). The yield and size distribution of the amplified libraries were assessed with Agilent Bioanalyzer using Ion AmpliSeq transcriptome Mouse gene expression Kit (Thermo Fisher Scientific). Sequencing was performed on an Ion Proton Instrument (Ion Torrent, Thermo Fisher Scientific). RNA-sequencing (RNA-seq) analysis was done using the Torrent Suite software version 5.4.0 and the RNASeqAnalysis plugin (Thermo Fisher Scientific) on the mouse reference genome mm10 using STAR. Gene level expression is calculated using HTSeq and Picard to report raw read counts and FPKM (fragment per kilobase of transcript per million).

### Cytolytic assay

EL4 cells (Target) were cultivated in RPMI + 10%FBS and incubated at RT for 25 min with Tag-it 1/1000.Then, EL4 cells were pulsed with 1 µg/ml of OVA peptide at 37°C for 1 hour. OT1 splenocytes (Effector) were isolated as previously described and activated with Anti-CD28 2ug/mL + IL-2 5ng/mL and anti-CD3e 2ug/mL in activation media for 1 day at 37°C. Effector cell were electroporated with 125nM of GZM siRNA or Ctrl siRNA and cultivated overnight in RPMI+20% FBS+2%L-Glutamine+anti-CD3 (2ug/mL) and anti-CD28 (2ug/mL) + IL2 (5ng/mL). Next day, cells were spin down at 300g for 5min, resuspended at the concentration of 10^6^/mL and cultivated for 24h in activated media + IL2 (50IU/mL). Next day 25000 of target cell were co-cultured for 6 days in activation media with 250000 effector electroporated cells; Ration (E:T)=10:1 and dead percentage of target cell (Tag-it +) were analysed in facs using a viability dye.

#### Bone marrow cultures with GM-CSF

Bone marrow cells, at a concentration of 10^6^ cells per well, were seeded into 6-well tissue culture plates, each containing 4 ml of medium consisting of RPMI 1640, supplemented with glutamine, penicillin, streptomycin, 2-mercaptoethanol (all from Invitrogen), 10% heat-inactivated fetal calf serum and GM-CSF at 20 ng/ml (Peprotech) according to Helft et al 2015 (102). This culture was incubated at various days to generate a dendritic cell enriched population. On day 2, 2 ml of the existing medium was removed and replaced with 2 ml of fresh media with GM-CSF. On day 3, all medium was replaced with 4 ml of fresh medium containing 20 ng/ml GM-CSF. In some experiments, IL-4 (Peprotech) was added to the medium at a concentration of 5 ng/ml, along with GM-CSF. On day 6-8, non-adherent cells from the culture supernatant and the loosely adherent cells were harvested by gentle washing with PBS, were collected and combined.

### Statistical analysis

Respect of the assumptions of normal distribution of residuals and homoscedasticity was verified, and data were presented as mean + SEM as well as individual values, unless otherwise indicated. Data from independent experiments were pooled for analysis in each data panel, unless otherwise indicated. Statistical analyses were performed using Prism 8 (GraphPad Software). Two-ways unpaired *t* test was performed unless otherwise indicated in the figure legends. We considered a P-value lower than 0,05 as significant. N.S.: *P*>0,05; *: *P*<0,05; **: *P*<0,01; ***: *P*<0,001; \*\*\*\**P*<0,0001.

## Data availability statement

Data sharing is not applicable as data for which a community-recognized, structured repository exist (e.g. next-generation sequencing data) were not created or analyzed in this study.

## Authors contribution

**Conception and design**: B.M., J.K. and C.E.R.

**Development of methodology**: B.M., J.K and A.K.

**Acquisition of data:** B.M., J.K, A.K., C.L, Y.G., N.A.P., L.C.P., L.H, A.L.C. and H.T.

**Analysis and interpretation of data:** B.M., J.K and A.K.

**Writing, review, and/or revision of the manuscript:** B.M. and C.E.R.

**Administrative, technical, or material support:** B.M., J.K, A.K. and C.E.R

**Study supervision:** B.M. and C.E.R.

---

**Fig. S1:**
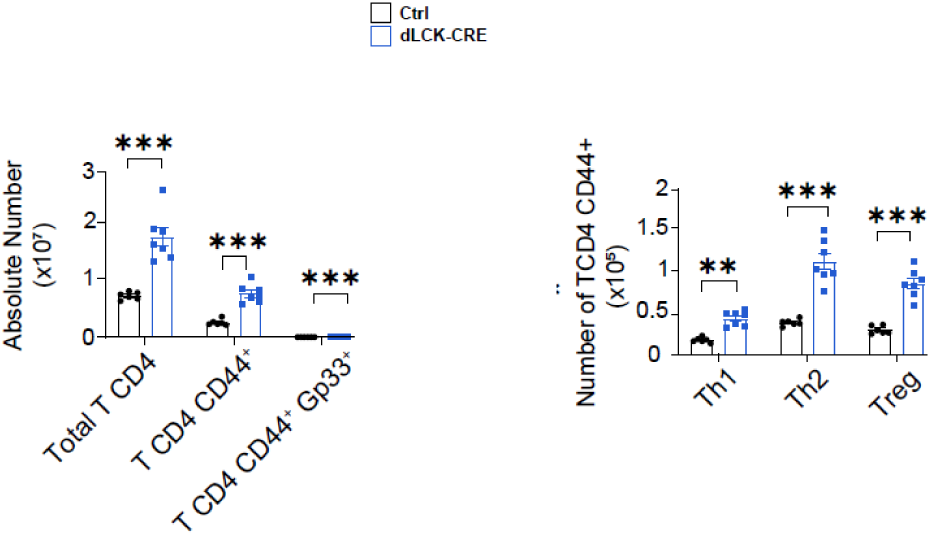
Flow cytometric characterization of splenic CD4+ T cell subsets in dLckCre (GSK-3KD) mice 30 days post-LCMV Cl13 infection. Left: Elevated numbers of total CD4 + T cells, CD44+CD4+ T cells, and CD44+Gp33+CD4+ T cells in GSK-3KD mice relative to controls. Right: Increased frequencies of Th1, Th2, and regulatory T cell populations in GSK-3KD mice. Analysis performed using anti-TCR beta, CD4, CD8, and FoxP3 antibodies.

**Fig. S2:**
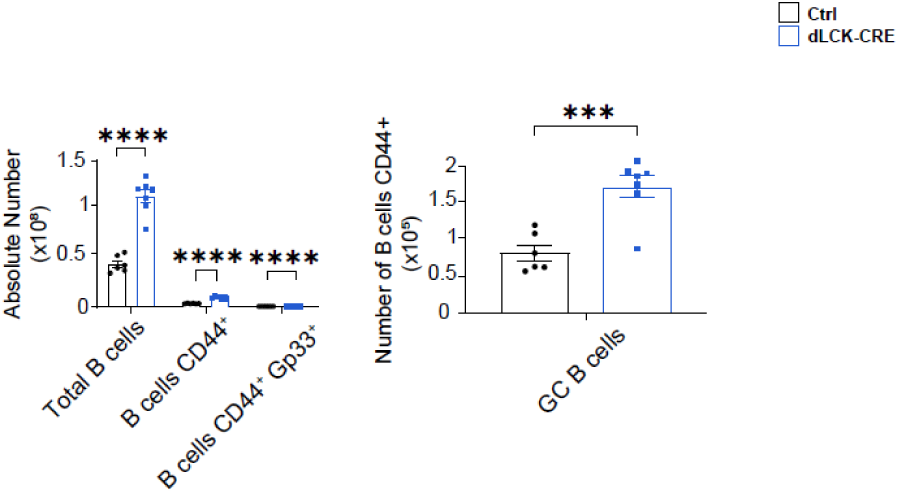
Flow cytometric analysis of splenic B cell populations in dLckCre (GSK-3KD) mice 30 days post-LCMV Cl13 infection. Left panel: Quantification of total B cells, CD44+ B cells, and CD44+Gp33+ B cells shows significant expansion in GSK-3KD mice compared to controls. Right panel: Enhanced germinal center CD44+ B cell numbers in GSK-3KD mice. Cell populations were identified using anti-CD19, CD44, and Gp33 tetramer staining.

**Fig. S3:**
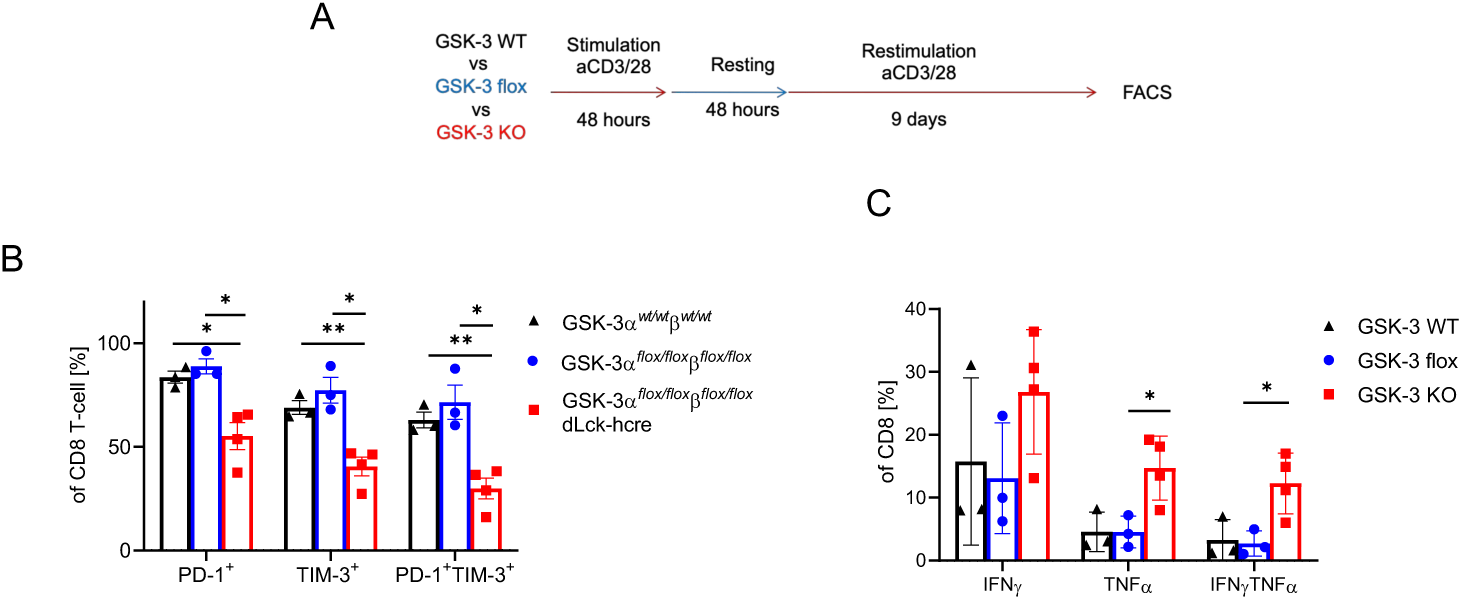
Knockout of GSK-3 in CD8 T cells confers resistance to *in vitro* exhaustion and enhances IFN-γ and TNF-α production. Splenic T cells were activated and restimulated according to the protocol outlined in Panel A. This process resulted in the upregulated expression of the exhaustion markers PD-1 and TIM-3 in both wild-type (WT) and floxed control T cells. Conversely, T cells deficient in GSK-3 (GSK-3-/- Lck Cre T cells) displayed significantly reduced levels of PD-1 and TIM-3 expression (B). Notably, the frequency of cells co-producing IFN-γ and TNF-α increased in GSK-3 Lck Cre T cells (C).

**Fig. S3:**
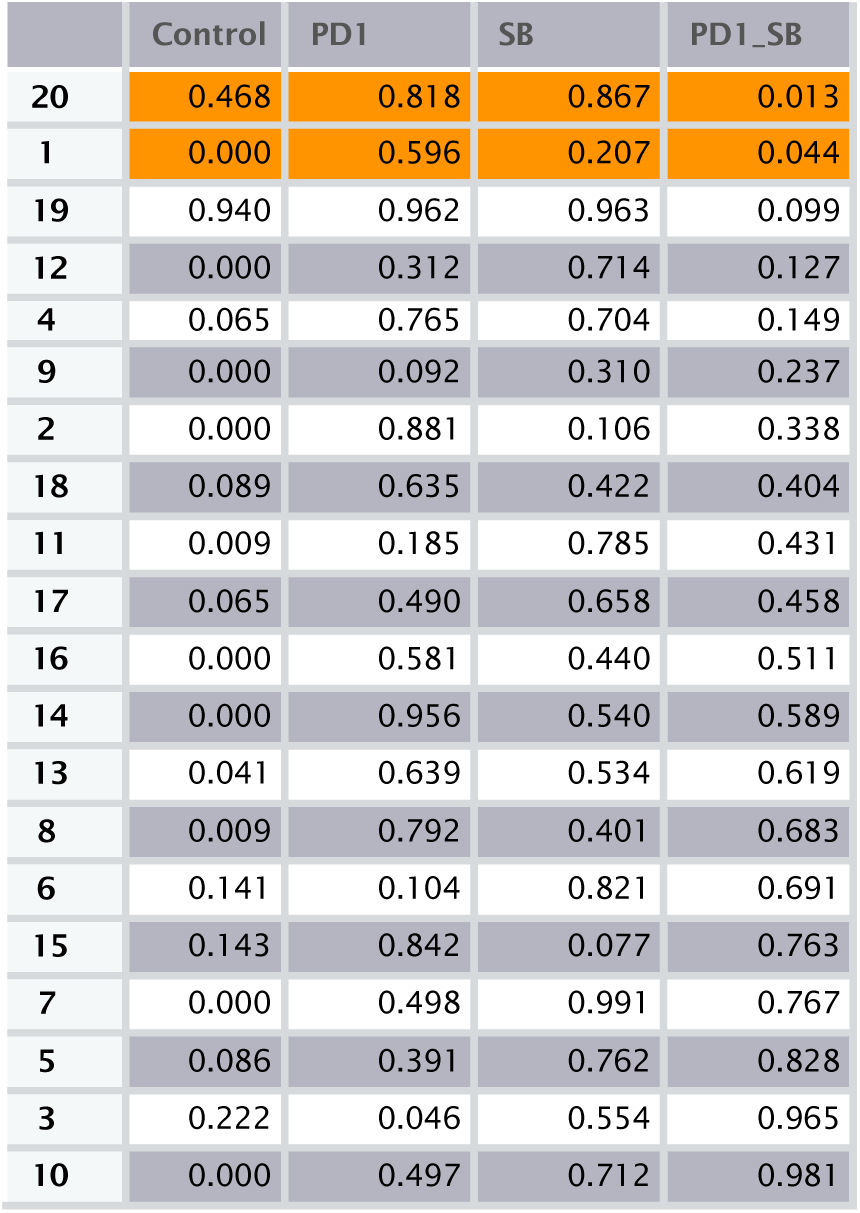
Statistical evaluation of mass cytometry (CyTOF) data comparing tumor-infiltrating lymphocyte subsets across treatment groups (control, anti-PD-1 monotherapy, SB415286 monotherapy, and combined anti-PD-1/SB415286). Clusters 20 and 1 showed statistically significant differences (p<0.05) for combination therapy relative to other treatment groups. Analysis based on three independent experiments with four mice per treatment group (n=3, 4 mice/group).

**Fig S5:**
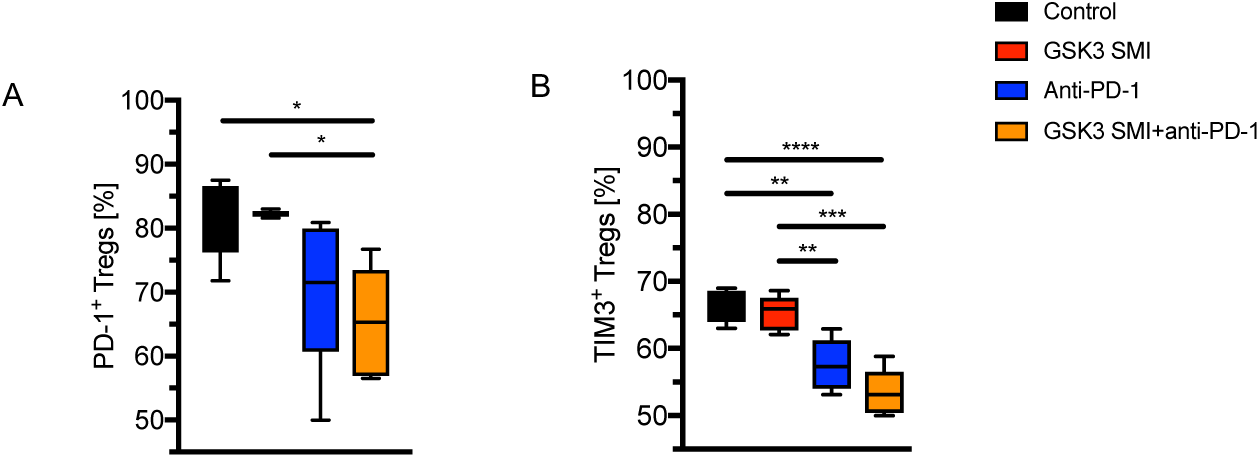
Flow cytometric validation showing the decrease in CD4+FoxP3+ TILs with reduced PD-1 and TIM3 expression in response to anti-PD1/GSK3i therapy. TILs were extracted from B16-F10 R1 tumors and stained for CD4, FoxP3, CD25, PD-1, TIM3, LAG3 and PDL-1 and assessed for expression. (A) Percentage of PD-1+ cells within the Treg population decreases with combination therapy. (B) Percentage of TIM3+ cells within the Treg population decreases with combination therapy.

**Figure S6:**
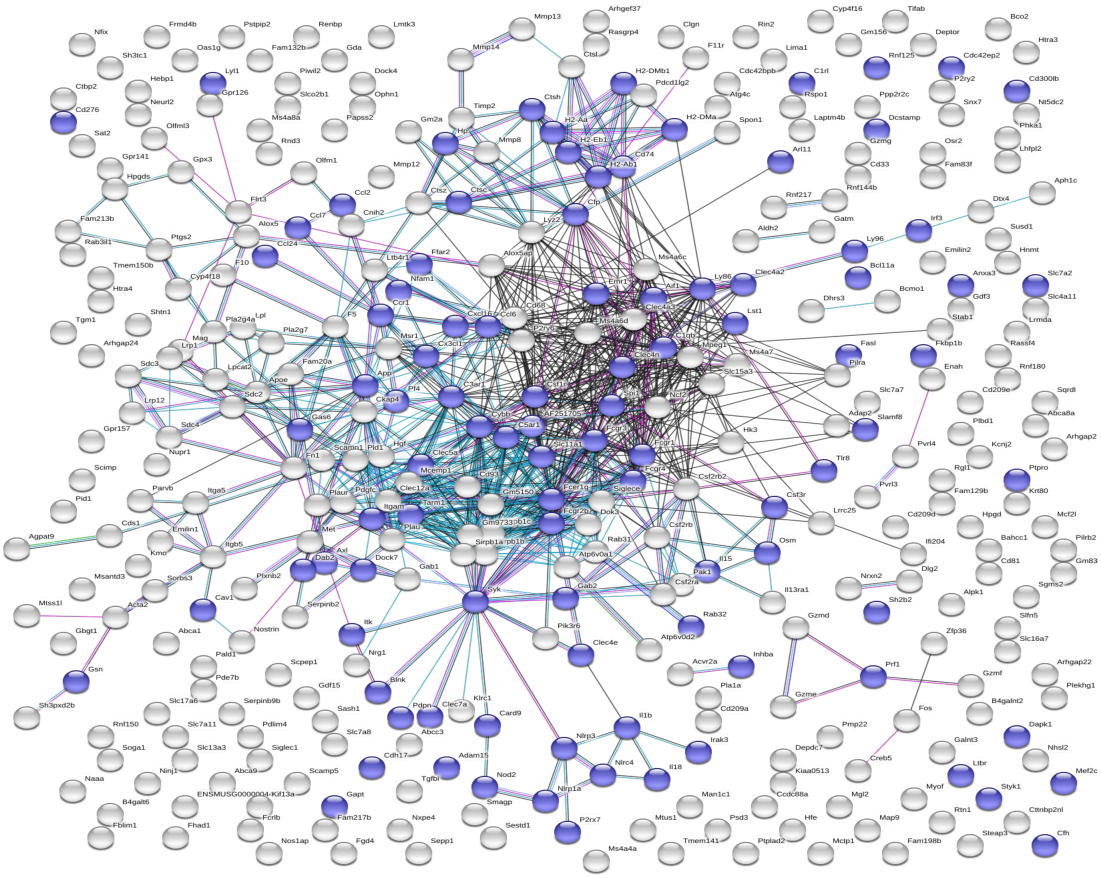
Gene regulatory network illustrating the synergistic upregulation of immunomodulatory gene signatures induced by combination treatment with anti-PD-1 and GSK-3 inhibition (see blue), as evidenced by the enrichment of genes associated with immune activation and inflammatory response pathways (shown as central network).

**Fig: S7.**
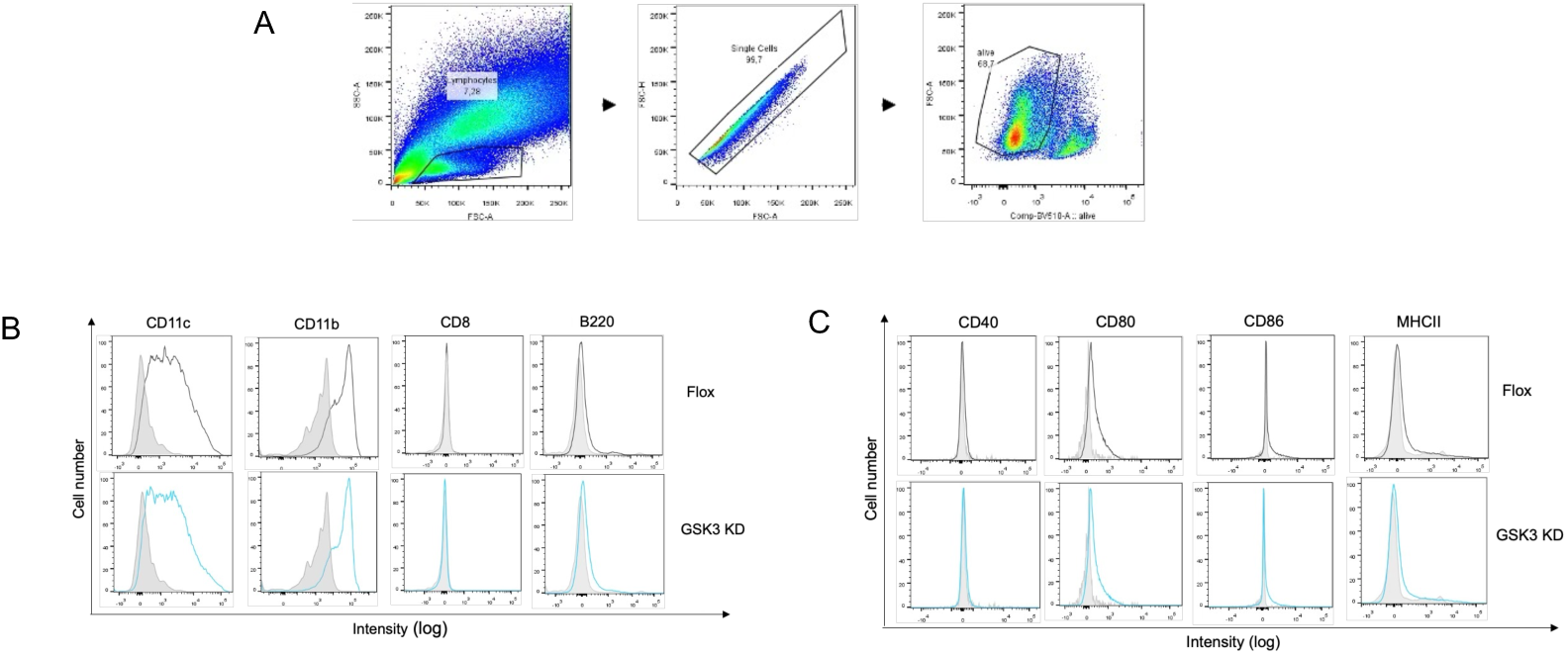
The phenotype of GM-CSF treated bone marrow dendritic cells from GSK-3KD and WT mice are the same. Bone marrow cells were isolated and cultured in medium containing GM-CSF (granulocyte-macrophage colony-stimulating factor), which stimulates the differentiation of hematopoietic progenitor cells into dendritic cells. Over the course of 6-8 days, the non-adherent cells are harvested and enriched for dendritic cells. (A) Gating strategy for DCs; (B) Flow cytometry panel showing equal expression of CD11c, CD11b and mutually low levels of CD8 and CD220. (C) Flow cytometry panel showing equal expression of CD80 and mutually low levels of MHC class II and CD86.

